# Functionally distinct ALK and ROS1 fusions detected in infant-type hemispheric gliomas converge on STAT3 and SHP2 activation

**DOI:** 10.1101/2025.05.27.656302

**Authors:** Andreas Postlmayr, Astrid Sanchez Bergman, Jacob Torrejon Diaz, Bernard Ciraulo, Shen Yan, Nina Hofmann, Samanta Carbajal, Rosalie Dobler, Charbel Machaalani, Laura Priego-Gonzalez, Andrea J. De Micheli, Ernesto Berenjeno-Correa, Luca Baroncini, Michael A. Grotzer, Uri Tabori, Cynthia Hawkins, Olivier Ayrault, Marc Zuckermann, Martin Baumgartner, Ana S. Guerreiro Stücklin

## Abstract

*ALK* and *ROS1* fusions are emerging as key drivers of infant-type hemispheric gliomas (IHG). With diverse gene partners, the impact of ALK and ROS1 oncoprotein heterogeneity on glioma biology remains unknown. We developed an integrative phospho-proteomic and transcriptomic approach to discover biological functions regulated by five IHG-associated fusions: *CCDC88A::ALK, PPP1CB::ALK, GOPC::ROS1, CLIP1::ROS1* and *KIF21A::ROS1*. Here, we report fusion-specific oncogenic functions conferred by the 5’ gene partner, including increased cell motility driven by microtubule-interacting fusions *CCDC88A::ALK* and *CLIP1::ROS1*. All studied fusions converged on a non-canonical STAT3 activation. Using affinity purification mass spectrometry, we identify SHP2 in direct interaction with all three ROS1 oncoproteins, but with none of the ALK oncoproteins, which in turn interact with SHC1/SHC3. ROS1 fusions phosphorylated SHP2 to a larger extent than ALK fusions, and analyses of downstream pathways suggested MAPK-independent, non-canonical SHP2-driven functions. Our findings reveal both common and fusion-specific dependencies, offering opportunities to optimize therapeutic strategies for pediatric gliomas.

## INTRODUCTION

Gene fusions are hallmarks of childhood cancers and are among the most common genetic drivers of human tumors (1,2). The spectrum of oncogenic fusions in pediatric central nervous system (CNS) tumors is unfolding (reviewed in 3), and recent studies have reported recurrent and novel gene fusions in up to 22.5% of tumors (2). Most fusions are detected in glial tumors, including pediatric low-grade gliomas (pLGG) (4), pediatric high-grade gliomas (pHGG) (5-7), and ependymomas (8). While some fusions are specific to tumor entities (e.g. *KIAA1549::BRAF* in pLGG), others are prevalent across many pediatric and adult cancers, and their roles in childhood brain tumor development remain to be elucidated.

Several gene fusions involve rearrangements between receptor tyrosine kinase (RTK) genes and multiple fusion partners, resulting in a vast range of activated oncoproteins. Recently, we and others reported a subset of gliomas prevalent in infants and young children — now referred to as *infant-type hemispheric gliomas* (IHG) —, which are driven by oncogenic fusions involving ALK, ROS1, NTRK1/2/3, and MET (henceforth collectively referred to as RTK fusions) (5,6,9). Multiple studies have previously reported the oncogenic properties of RTK-fusions in pediatric and adult cancers, including non-small cell lung cancer (NSCLC), anaplastic large cell lymphoma (ALCL), colorectal cancer and inflammatory myofibroblastic tumors (reviewed in 10). In gliomas, ALK fusions appear to occur predominantly in IHGs, whereas ROS1 fusions were initially reported in adult gliomas (11); NTRK family kinase fusions have been detected in several pLGG and pHGG (7) and MET fusions predominantly in other pHGGs (9,12).

In healthy tissues, these RTKs are membrane-bound and activated upon ligand binding, thereby orchestrating numerous cellular processes, including growth, proliferation, migration, differentiation, and apoptosis, via RAS/MAPK, PI3K/AKT/mTOR, and/or JAK/STAT signaling cascades, among others. RTK fusion gene breakpoints are relatively conserved, typically leading to retention of the C-terminal region containing the kinase domain and loss of the extracellular and transmembrane domains. There are multiple ALK and ROS1 inhibitors in clinical use and initial reports demonstrate favorable responses to Lorlatinib in ALK-driven IHGs and to Entrectinib in ROS1-driven IHGs (13,14). However, both tumor progression on treatment and tumor recurrence upon treatment discontinuation have been observed, raising concerns regarding the long-term efficacy of RTK inhibitors for children with IHGs, especially when used in monotherapy (13,14).

Although the oncogenic properties of RTK fusions have been demonstrated in diverse human cancers, the molecular mechanisms underlying individual fusions within this broad group of alterations remain largely unknown. Focusing on RTK fusions enriched in IHGs and using an integrated multi-omics approach, we explored the mechanistic roles of five distinct ALK- and ROS1-fusions, uncovering specific pathway and phenotypic dependencies. Here we report SHP2 as a signaling effector activated and present in direct interaction with ROS1 fusions, whereas the adaptor proteins SHC1/SHC3 were detected in protein complexes with ALK fusions. We identified direct, non-canonical STAT3 activation downstream all studied ALK- and ROS1-fusions. We further uncovered and characterized fusion-specific oncogenic phenotypes, particularly the role of fusions with microtubule-interacting fusion partners in cell motility. Our findings provide mechanistic insights and molecular information that can be leveraged to develop tailored combinatorial treatment approaches for ALK- and ROS1-fusion driven IHGs.

## RESULTS

### ALK, ROS1, MET and NTRK fusions are prevalent across a diverse spectrum of pediatric gliomas, with distinct co-alterations and age distribution

ALK, ROS1, MET or NTRK family fusions are a hallmark of IHGs but not exclusive to this tumor entity. To obtain a comprehensive view of their distribution across pediatric gliomas, we analyzed data from the OpenPedCan (version 15) project. Data of 1524 glioma patients were deposited, and upon manual filtering for potentially false positive fusion annotations, 8 ALK, 7 ROS1, 19 MET and 25 NTRK family fusion harboring patients were identified, which corresponds to 66 deposited sample data and 3.87% of pediatric gliomas from the age of 0 to 21 years (Figure 1A).

**Figure 1:**
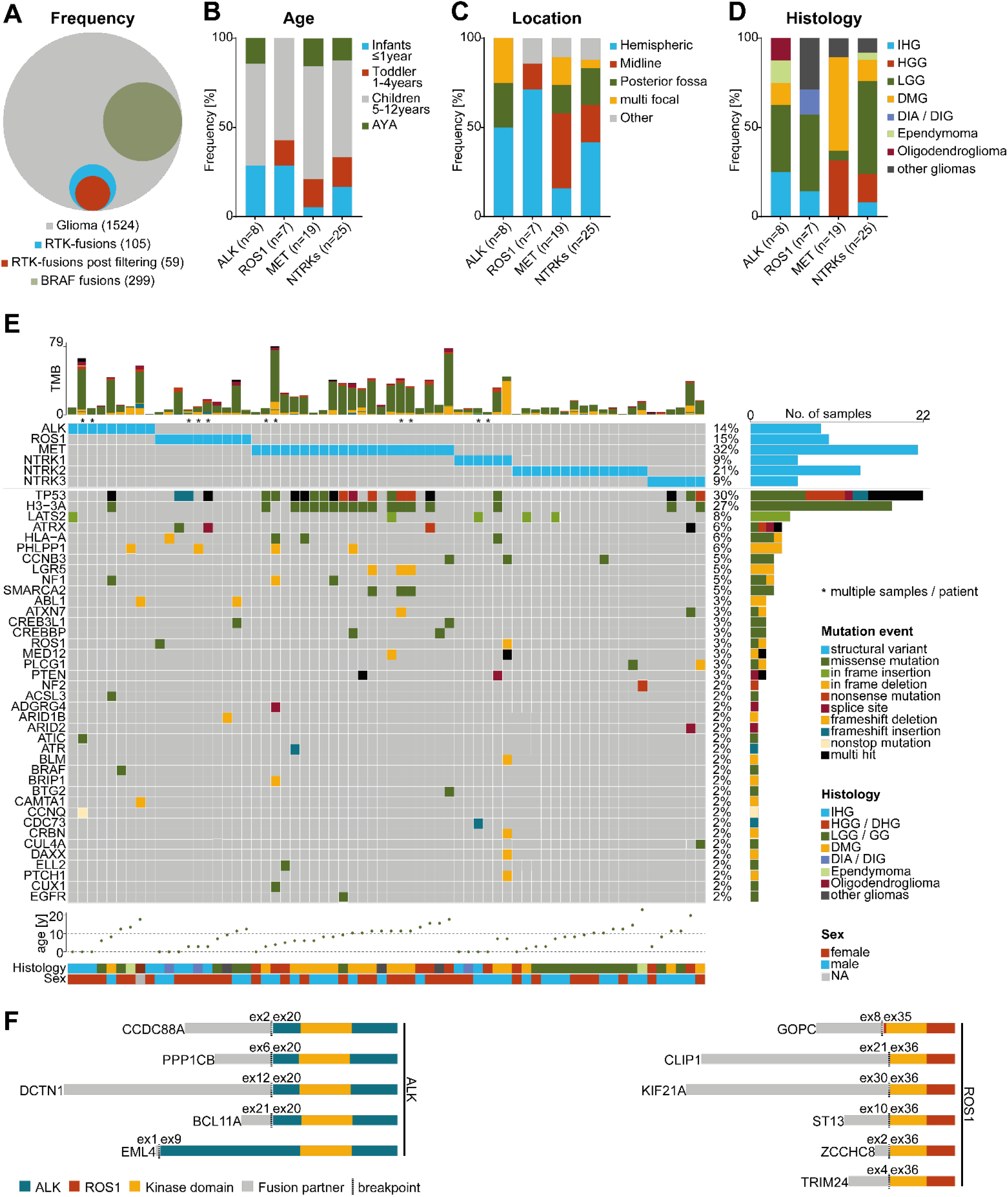
RTK-fusions are identified in pediatric CNS glioma entities across all ages and localizations. **(A)** Frequency of RTK-fused tumors in analyzed cohort. **(B-D**) Age **(B)**,localization **(C)**, and histology **(D)** distribution of RTK-fusion harboring gliomas; x-axis: ALK-fusions (n=8), ROS1-fusions (n=7), MET-fusions (n=19), NTRK-fusions (n=25); y-axis: distribution in percentage. **(B)** blue: infants 0<1 years, red: toddlers 1-4 years, grey: children 5-12 years, green: adolescent and young adults (AYA) >13 years. **(C)** blue: hemispheres, red: midline, green: posterior fossa, yellow: multi focal, grey: other. **(D)** blue: infant-type hemispheric gliomas (IHG), red: high grade gliomas (HGG) and diffuse hemispheric gliomas (DHG), green: low grade gliomas (LGG) and ganglioglioma (GG), yellow: diffuse midline glioma (DMG), purple: diffuse infantile astrocytoma (DIA) and desmoplastic infantile ganglioglioma (DIG), light green: ependymoma, berry: oligodendroglioma; petrol: other gliomas. **(E)** oncoprint indicating the top 40 co-occurring oncogenic mutations in RTK-fusion harboring gliomas; top bar-plot: tumor mutational burden (TMB), detected mutations per megabase; matrix: indicating type of mutation detected per RTK-fusion; right bar-plot: frequency of alterations by gene; alterations: blue: structural variant, dark green: missense mutation, light green: in frame insertion, yellow: in frame deletion, orange: nonsense mutation, berry: splice site, grey: frameshift deletion, petrol: frameshift insertion, black: multi hit; bottom scatter plot: age of patients in years; bottom graphs: upper: histology: characterization according to database: blue: infant-type hemispheric gliomas (IHG), red: high grade gliomas (HGG) and diffuse hemispheric gliomas (DHG), green: low grade gliomas (LGG) and ganglioglioma (GG), yellow: diffuse midline glioma (DMG), purple: diffuse infantile astrocytoma (DIA) and desmoplastic infantile ganglioglioma (DIG), light green: ependymoma, berry: oligodendroglioma; petrol: other gliomas; lower: sex, blue: male, red: female; *: multiple samples from the same patient. **(F)** Schematic representation of ALK- and ROS1 fusions identified in cohort; left: ALK-fusions, right: ROS1-fusions; blue and red: C-terminal part of ALK or ROS1 maintained in fusion, grey: N-terminal part of fusion partner protein, yellow: kinase domain, horizontal black line: fusion breakpoint, ex: exons fused in fusion protein, schematics scaled according to protein size.

ALK- and ROS1-fusions showed a higher prevalence in infants (each at 28.6%) and localization to hemispheres (50% of ALK-, and 71.4% of ROS1-fusions) than in MET- and NTRK-fusions (infants: MET-fusions 5.3%, NTRK-fusions: 16.7%; hemispheric: MET-fusions: 15.8%, NTRK-fusions: 30.3%) (Figure 1B,C). Histopathological annotation varied, with ALK-, ROS1-, and NTRK-fused gliomas spanning a broad spectrum of gliomas and other tumor entities. However, this was not centrally reviewed or adjusted according to the current WHO classification of CNS tumors. Most ALK-fused tumors were labeled as low-grade gliomas or gangliogliomas (LGG / GG) (37.5%) or infant-type hemispheric gliomas (IHG) (25%).

ROS1-fusions were detected in tumors classified as LGG / GG (42.9%), followed by other rare CNS tumor histologies (28.6%) and IHGs (14.3%). MET-fused tumors were described as diffuse midline gliomas (DMG) (52.6%), and high-grade and diffuse-hemispheric gliomas (HGG / DHG) (31.6%). LGG / GG (52%) was the most prominent histology for NTRK-fusions (Figure 1D).

Genomic characterization of these tumors further revealed that 36% of tumor samples did not harbor additional mutations in known oncogenes or tumor suppressor genes. Most additional recurrent alterations were detected at *TP53* (30%) or *H3F3A* (24%) level, both enriched in MET-fused samples (63%). The presence of the H3 K27M mutation (11 of 19 MET-fused tumors) was consistent with a DMG diagnosis (Figure 1E).

Further analyses of ALK- and ROS1-fusions revealed the presence of conserved breakpoints (except for *EML4::ALK*) (Figure 1F). These fusion partners include a group of cytoskeleton- and microtubule-associated proteins, CCDC88A, DCTN1, EML4, CLIP1, and KIF21A, which are known to be involved in cell motility, mitotic spindle assembly, and intracellular transport. Other fusion partners, such as BCL11A, a transcription factor involved in neuronal development, and TRIM24, an E3 ubiquitin ligase involved in chromatin remodeling, are nuclear proteins. In the third group, proteins with very specific subcellular localization and function were also detected as RTK fusion partners: GOPC, a Golgi apparatus-associated protein; ST13, a chaperone for heat-shock proteins; PPP1CB, a phosphatase involved in cell division; and ZCCHC8, involved in RNA processing.

Detailed characterization of this patient cohort highlights that ALK-, ROS1-, MET-, and NTRK-fusions occur across several pediatric CNS tumor entities. The broad spectrum of fusion partners observed aligns with previous reports, raising the question of whether heterogeneous fusions can differentially impact cellular functions and disease outcomes. Finally, the paucity of additional oncogenic alterations in most ALK and ROS1-fused tumors further corroborates the role of RTK fusions as the main oncogenic tumor drivers in these tumors.

### Kinase function is indispensable, and fusion partners are required for the tumorigenic potential of ALK- and ROS1-fusions

Next, we sought to understand whether kinase function and fusion partners are essential for tumorigenic behavior. To this end, we generated cell models of two ALK-fusions (CCDC88A::ALK and PPP1CB::ALK) and three ROS1-fusions (GOPC::ROS1, CLIP1::ROS1, and KIF21A::ROS1). We mutated the ATP binding site within the kinase domain to generate kinase dead mutant (KD) variants of the fusion genes, which we used as controls (Figure 2A). We used immortalized normal human astrocytes (iNHAs) and transgenically introduced fusion constructs or their respective KD control variants. Western blot analysis confirmed the expression of all fusions and phosphorylation of ALK (Tyr1507) and ROS1 (Tyr2274), which were absent in the KD controls (Figure 2B).

**Figure 2:**
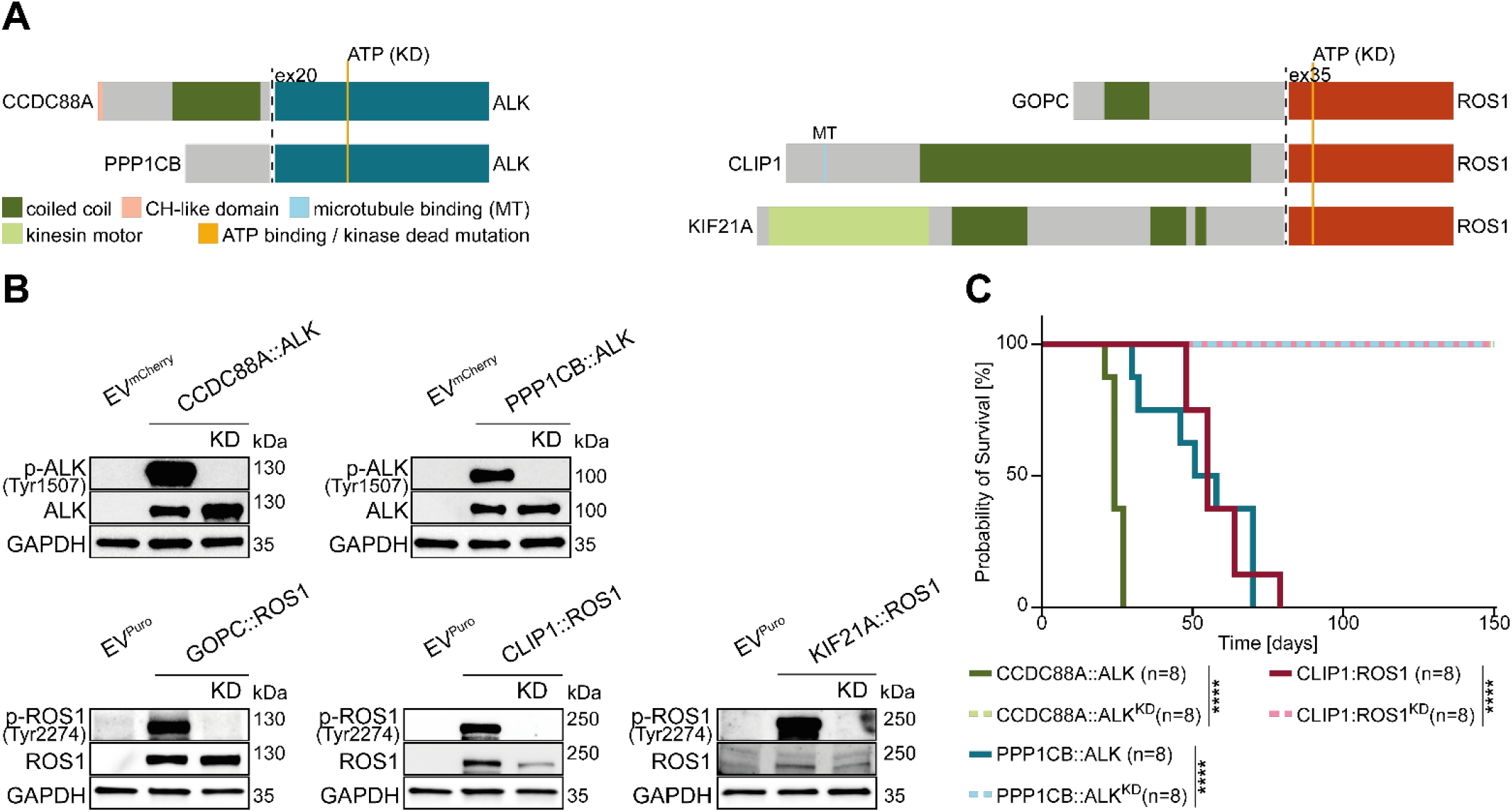
Intact kinase domain is indispensable for ALK and ROS1-fusion tumorigenesis. **(A)** Detailed schematics of ALK-(left) and ROS1-fusions (right) used in the study; petrol and red: C-terminal part of ALK or ROS1 maintained in fusion, grey: N-terminal part of fusion partner protein, yellow: ATP binding site of ALK or ROS1 altered to generate kinase dead mutants, fusion partner protein domains: green: coiled coil domain, pink: CH domain, light blue: microtubule binding, dark grey: active site, light green: kinesin motor domain, horizontal black line: fusion breakpoint, ex20 (left), ex35 (right): first ALK or ROS1 exon, respectively, maintained in fusion protein; schematics scaled according to protein size. **(B)** Western blot validation of ALK- and ROS1-fusion transgene expression in iNHAs, schematically set out in (A); p-ALK (Tyr1507): ALK activating phospho site, p-ROS1 (Tyr2274): ROS1 activating phospho site, GAPDH: loading control, phospho specific antibodies: validation of kinase dead mutation, EV^mCherry/Puro^: empty vector control, KD: kinase dead construct. **(C)** Kaplan-Meier survival curves showing tumor induced mortality upon orthotopic intracranial injection of mutant iNHAs in NSG mice, groups are represented by individual curves, with a n=8 mice per group, solid lines functional ALK-or ROS1-fusions, dashed lines: corresponding KD controls; green: CCDC88A::ALK, petrol: PPP1CB::ALK, berry: CLIP1::ROS1; log-rank test, ****: p-value<0.0001

To test their role in cell transformation and tumor formation, we intracranially injected our cell models into NOD scid gamma (NSG) mice. All fusion transgenes resulted in tumor formation in all mice injected.

The median survival was 24 days for CCDC88A::ALK, 54.5 days for PPP1CB::ALK, and 55 days for CLIP1::ROS1, compared to their respective KD controls, which did not result in tumor formation (Figure 2C). These orthotopic models validated the tumorigenicity of ALK and ROS1 fusions, confirming that a functional kinase domain is essential and that fusion partners contribute to oncofusion-induced tumorigenesis.

### Integrated (phospho-)proteomics and transcriptomics reveal fusion-specific cell signaling and pathway dependencies

We investigated fusion-specific regulation of gene expression and protein activation by bulk mRNA sequencing and (phospho-)proteomic mass spectrometry, including phospho-tyrosine enrichment. We compared ALK / ROS1 fusions and their KD controls, both expanded *in vitro* due to the lack of *in vivo* engraftment of KD controls (Figure 2C). Unbiased hierarchical clustering of the most differentially regulated molecules detected in each omics layer revealed that distinct alterations were primarily driven by the respective RTK (Figure S1A). These results led to the separation of ALK- and ROS1-fusion models for subsequent analyses.

To further explore the molecular complexity of ALK- and ROS1-fusion models, we integrated transcriptome, proteome and phosphoproteome data using an unsupervised joint dimensional reduction (jDR) approach called Multi-Omics Factor Analysis (MOFA) (15). This integrative approach uncovered latent factors explaining biological variability that was not detectable through analysis of individual omics layers (Figure 3A).

**Figure 3:**
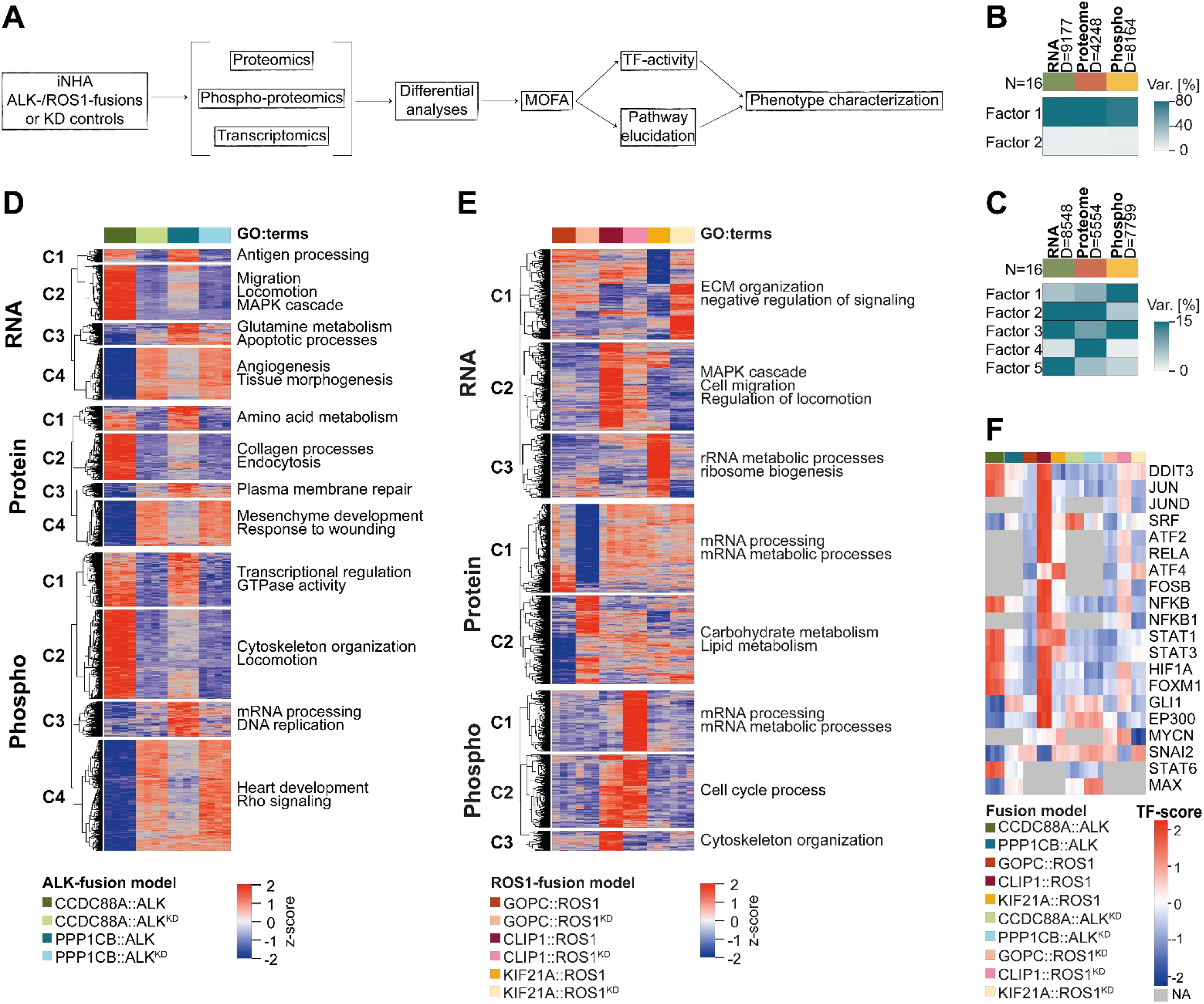
Integrative phosphoproteotranscriptomic analyses reveal upregulated STAT and MAPK activity and distinct dysregulation of cellular functions in ALK- and ROS1-fusions. **(A)** Flowchart outlying multi-omics approach. **(B**,**C)** MOFA variance contribution matrix; top bar: omic layer set size, green: transcriptomics, red: proteomics, yellow; phosphoproteomics, D: events used for analysis, N: sample size; heatmap: proportion of variance each latent factor accounts for across omic layers in percent, **(B)** ALK-fusion samples, **(C)** ROS-fusion samples. **(D**,**E)** MOFA heatmap illustrating factor-specific variance contributions for **(D)** ALK- and **(E)** ROS1-fusions; clusters based on mutant iNHA models; GO::terms: enriched pathways based on factor relevant molecules; color gradient: z-score high (red) to low (blue); **(E)** focused heatmap from FigS3A. **(F)** Focused heatmap illustrating transcription factor activity based on MOFA weights; color gradient: z-score high (red) to low (blue), grey: NA.

For ALK-fusions, two latent factors were identified, accounting for up to 80% of the variance across all data sets (Figure 3B, Figure S1B). In contrast, MOFA of ROS1-fusions revealed five latent factors, explaining up to 15% of the observed variance (Figure 3C), reflecting the heterogeneity among the three ROS1-fusions models (Figure 3B,C, Figure S1B).

Next, we performed unsupervised hierarchical clustering of the most variable features. In ALK-fusions, four distinct clusters consistently emerged across all three omics layers, specifically characterizing CCDC88A::ALK, PPP1CB::ALK, both fusions, or both kinase-dead (KD) controls (Figure 3D). ROS1-fusions displayed more heterogeneous clustering, with up to eight clusters per omics layer and limited concordance across layers, further supporting broader differences among ROS1-fusions (Figure S2A). A focused unsupervised hierarchical clustering approach, using only features with the strongest contribution to specific MOFA factors (Figure 3C, Figure S1B) resulted in a reduced number of clusters (Figure 3E).

Functional enrichment analysis using Gene Ontology (GO) indicated specific cellular programs regulated by the fusion proteins. ALK-fusions were associated with an upregulation of antigen-presenting signatures, metabolic programs, as well as DNA replication and transcriptional regulation, which can be indicative of proliferative cell states (Figure 3D). Interestingly, several developmental programs were repressed in ALK-fusions, including tissue morphogenesis and angiogenesis, mesenchymal and heart development. This repressive pattern might be representative of a broader block in development driven by ALK-fusions. ROS1 fusions displayed a fusion-specific landscape of deregulated biological functions. For the Golgi associated GOPC::ROS1, we found an enrichment for pathways involved in mRNA processing and mRNA metabolic processes, whereas for KIF21A::ROS1, an increased rRNA metabolism signature was detected (Figure 3E). Notably, for both cytoskeleton-associated fusion proteins CLIP1::ROS1 and CCDC88A::ALK we identified an upregulation of signatures related to locomotion, cytoskeletal organization and MAPK signaling across various omics layers (Figure 3D,E).

We next assessed transcription factor (TF) activities (Figure 3F, S2B). ALK-fusions displayed upregulation of TFs involved in stemness (e.g.: STAT3 and HIF1α) and proliferation (e.g.: FOXM1 and GLI1). ROS1-fusions, in contrast, showed deregulation of TFs involved in tissue differentiation (e.g.: EP300 and MYCN) and epithelial-to-mesenchymal transition (EMT) (e.g.: JUN and SNAI2). STAT3 increased transcriptional activity was also identified in ROS1-fusions, highlighting its importance in maintaining cellular plasticity in ALK- and ROS1-fusion cells. Furthermore, effectors of the MAPK signaling cascade (e.g.: FOS and JUN) were identified, more pronouncedly in ROS1-fusions. Finally, to further investigate the differences observed in MAPK signaling, we analyzed the phosphorylation levels of all MAPK signaling proteins annotated in KEGG (hsa04010). CCDC88A::ALK and CLIP1::ROS1 displayed increased phosphorylation levels in several, yet distinct, MAPK pathway members, whereas the other three fusions appeared to impact MAPK pathway protein phosphorylation to a lesser extent (Figure S2C). Collectively, integrative multi-omics analyses revealed both common and fusion-specific regulation of cellular functions and intracellular pathway activity.

### CCDC88A::ALK and CLIP1::ROS1 show increased invasive and motile phenotypes

Integrative multi-omics analyses revealed dysregulation in cell migration, locomotion and cytoskeleton organization, specifically in iNHA expressing CCDC88A::ALK and CLIP1::ROS1 (Figure 3D and E, Figure S3A). Notably, CCDC88A and CLIP1 (as well as KIF21A) contain microtubule (MT)-binding domains. To investigate the relevance of the microtubule interaction domains in those fusions, we abolished the MT-binding domain of CLIP1::ROS1 through substitution mutation (K98E and N99D; CLIP1::ROS1^ΔTUB^). Western blot analysis confirmed unaltered p-SHP2 and p-ERK1/2 levels in CLIP1::ROS1^ΔTUB^ but revealed reduced levels of p-STAT1/3 compared to functional CLIP1::ROS1 (Figure 4A). Phenotypic elucidation by immunofluorescence microscopy revealed morphological changes in CCDC88A::ALK and CLIP1::ROS1 cells, displaying an elongated, polarized cell shape compared to controls (EV; EV^mCherry^ for ALK-fusions and EV^Puro^ for ROS1-fusions, respectively) (Figure S3B). Notably, CLIP1::ROS1^ΔTUB^ showed different morphology compared to non-mutated CLIP1::ROS1, resembling the morphology of controls (Figure S3B). This observation indicates that a functional MT-binding domain impacts cell morphology. To phenotypically assess the MOFA postulated altered motility we evaluated the motile behavior and speed of these cells by live cell tracking imaging analyses (Figure 4B,C). Indeed, live cell tracking quantification revealed significantly increased cell speed in CCDC88A::ALK and CLIP1::ROS1 expressing cells compared to controls (Figure 4B). Notably, abolishing the MT domain resulted in reduced motility (Figure 4B).

**Figure 4:**
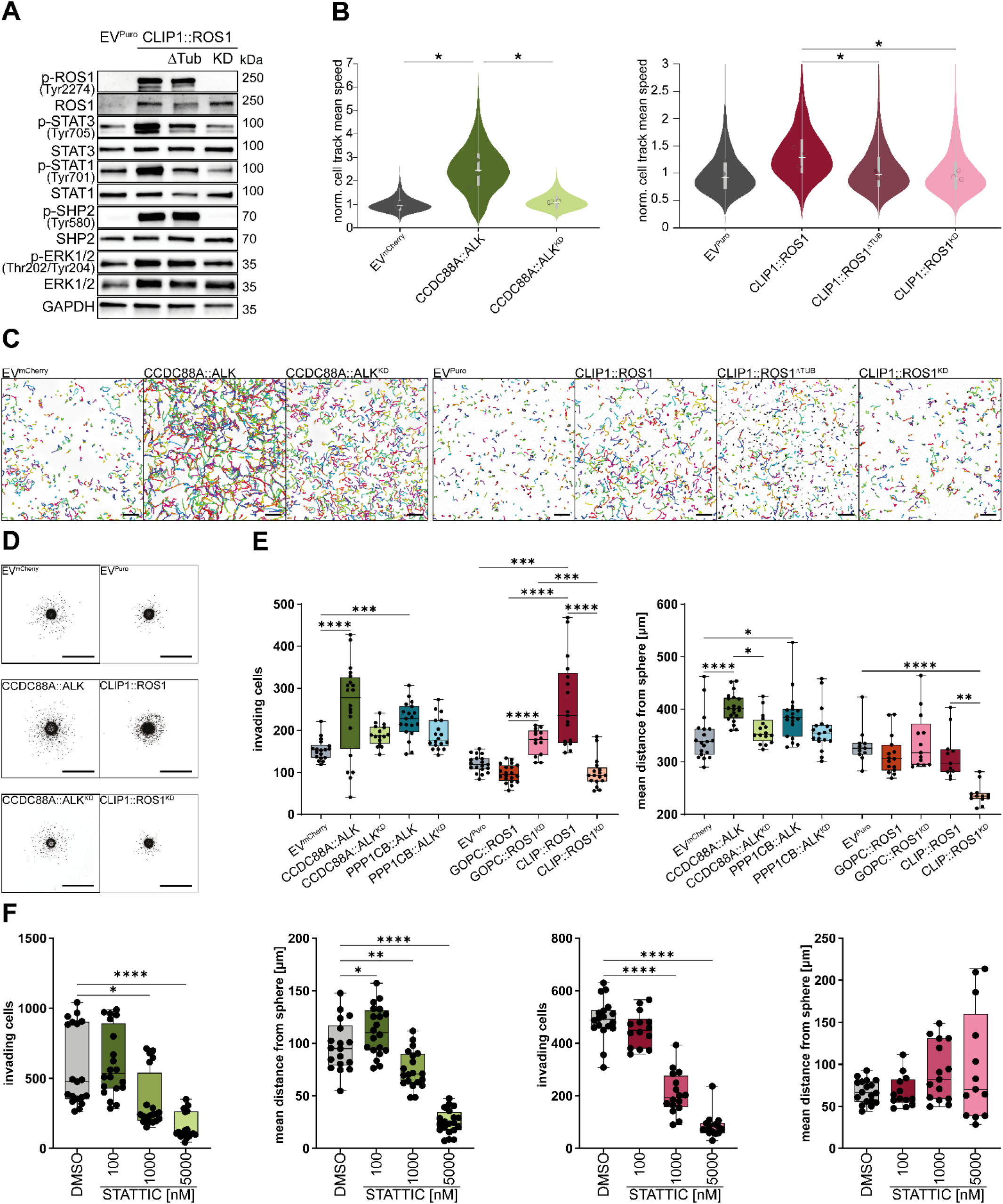
CCDC88A::ALK and CLIP::ROS1 expression result in hyper motile phenotypes. **(A)** Western blot analysis of CLIP1::ROS1-fusion expression and SHP2/MAPK and STAT3 signaling in CLIP1::ROS1-fusion iNHA; ΔTUB: abrogated microtubule interaction domain; GAPDH: loading control, p-ROS1 (Tyr2274) antibody used to validate fusion transgene activity, phospho-SHP2 (Tyr580), and p-ERK1/2 (Thr202/Tyr204) used to validate MAPK pathway activity and p-STAT3 (Tyr705), and p-STAT1 (Tyr701) for STAT activation. **(B)** Violin plots highlighting track mean speed from (C) for CCDC88A::ALK (left) and CLIP1::ROS1 (right); dots represent mean of biological replicates, significance calculated on mean values, significance calculated using unpaired two-tailed Student’s t-test, *: p-value ≤0.05. **(C)** Illustrative images of live cell tracking; inverted nuclear fluorescence, colored lines visualize tracks of individual cells within 12 hours; scalebar: 200µm. **(D)** Illustrative images of SIA assays; scalebar: 500µm. **(E)** SIA quantification of invading ALK- and ROS1-fusion iNHAs; left graph: number of invading cells, right graph: mean distance of invasion; one-way ANOVA (normally distributed) or Kruskal Wallis test (not normally distributed), post-hoc Dunn-Bonferroni test, *: p-value≤0.05, **: p-value <0.01,***:p-value<0.001, ****: p-value <0.0001. **(F)** SIA quantification of invading CCDC88A::ALK (first two graphs) or CLIP1::ROS1 (last two graphs) iNHAs treated with indicated STAT3i (Stattic) concentrations; first and third graph: number of invading cells, second and fourth graph: mean distance of invasion; one-way ANOVA (normally distributed) or Kruskal Wallis test (not normally distributed), post-hoc Dunn-Bonferroni test, *: p-value≤0.05, **: p-value <0.01, ****: p-value <0.0001.

Finally, we assessed the invasive properties of these fusions using spheroid invasion assays (SIA). CCDC88A::ALK- and CLIP1::ROS1-expressing cells showed a significant increase in number of invading cells, as well as an increase in the mean distance of invasion for CCDC88A::ALK cells compared to controls, and compared to PPP1CB::ALK, which also led to a modest increase in invasion (Figure 4D,E). Similarly, CLIP1::ROS1-expressing cells showed a significant increase in invasive behavior compared to the controls and to GOPC::ROS1. Invasion of CLIP1::ROS1 cells resembles a collective pattern, whereas CCDC88A::ALK cells appear to invade the surrounding matrix individually. These differences suggest distinct effects of CCDC88A::ALK and CLIP1::ROS1 on cytoskeletal rearrangements and invasion-promoting pathways.

Finally, given the reduction of p-STAT3 observed in CLIP1::ROS1^ΔTUB^ mutant cells, we tested the impact of a STAT3 inhibitor (Stattic) in CCDC88A::ALK- and CLIP1::ROS1-associated invasion. Indeed, the number of invading cells was dose-dependently reduced in both models. CCDC88A::ALK cells also showed a reduced mean distance of invasion when treated with Stattic (Figure 4F, Figure S3C).

Taken together, these data confirmed an increase in invasive and motile phenotypes in cells expressing fusions with MT-interacting domains. Further, our data suggest that cell invasion and motility are regulated by STAT3 signaling.

### ALK and ROS1 fusions interact with different signal transducing proteins and exhibit heterogenous levels of activation of MAPK signaling

Integrative multi-omics analyses revealed dysregulated MAPK signaling, especially in CCDC88A::ALK and CLIP1::ROS1 expressing cells. Next, we used affinity purification mass spectrometry (AP-MS) to identify direct interactors of the five studied fusions. We identified SHC1/3, known RTK adaptor proteins, as direct interactors of ALK-fusions and *PTPN11*/SHP2 as a direct interactor for all three ROS1-fusions (Figure 5A). We confirmed the direct interaction of CCDC88A::ALK with SHC1 and SHC3 by co-immunoprecipitation, with SHC1 interacting in its phosphorylated, i.e. active, form and the direct interaction of PPP1CB::ALK with SHC1 (Figure 5B). Further, we could validate the ROS1-fusions interaction with phosphorylated SHP2 (Figure 5B). Finally, using *in vitro* kinase assays we validated SHP2, at position Tyr580, as a direct substrate of ALK- and ROS1-fusions, irrespective of direct or indirect interaction (Figure S4A).

**Figure 5:**
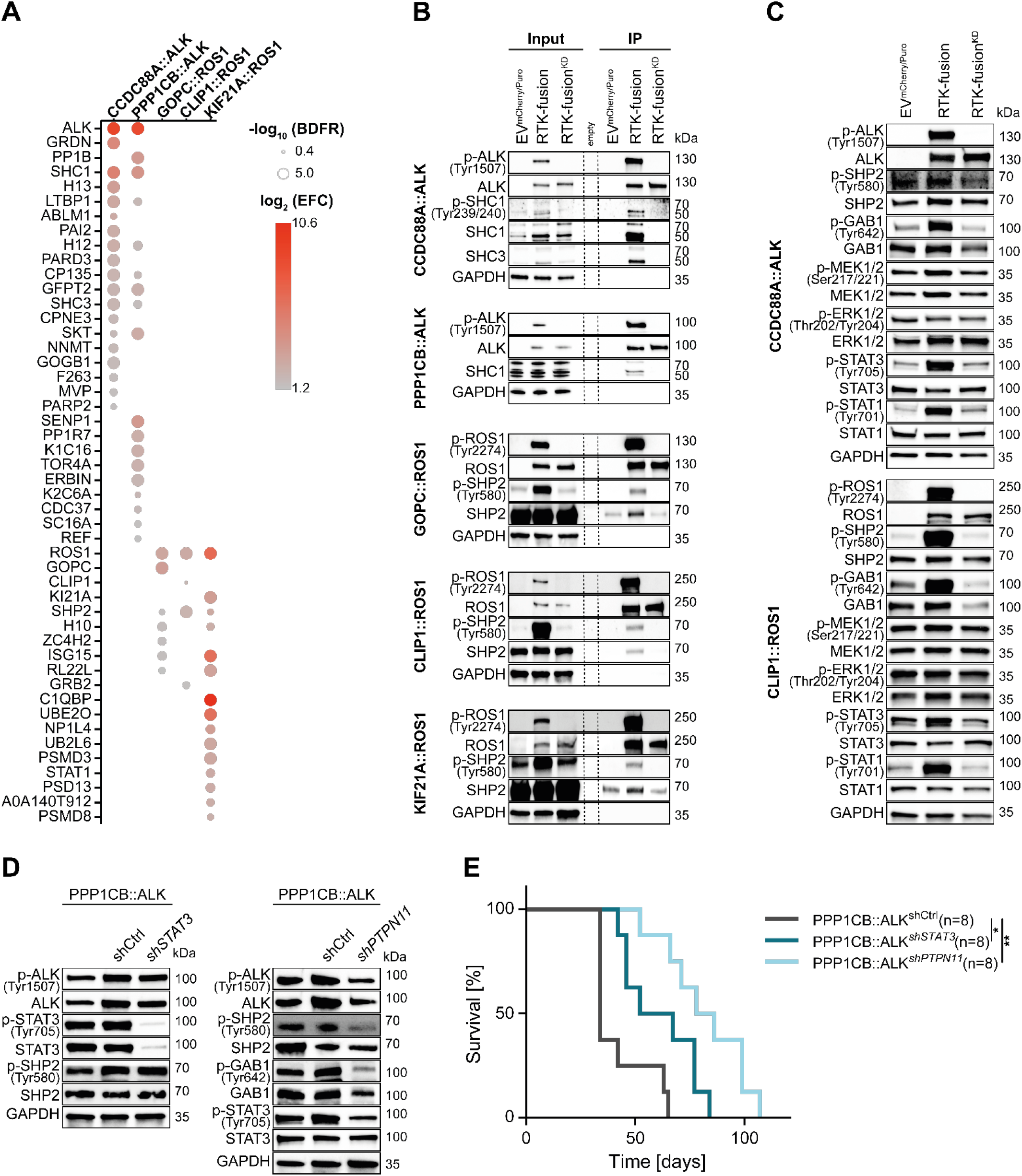
ALK- and ROS1-fusions converge on STAT3 and SHP2 activation. **(A)** Affinity purification MS/MS identifying direct interactors of ALK- and ROS1-fusions used in this study; size: −log_10_ BFDR, color gradient: log_2_ EFC high (red) to low (grey). **(B)** Immunoprecipitation validating SHC1/3 as direct interactors of ALK-fusion (top two blot) and SHP2 as direct interactor of ROS1-fusion (bottom three blots),respectively; GAPDH: loading control, p-ALK (Tyr1507), -ROS1 (Tyr2274) antibody used to validate KD mutants, p-SHC1 (Tyr239/240), and p-SHP2 (Tyr580) antibodies used to validate activity of interactors; dashed lines: marker lane. **(C)** Western blot analysis of MAPK signaling in CCDC88::ALK and CLIP1::ROS1 models. GAPDH: loading control, p-ALK (Tyr1507), -ROS1 (Tyr2274) antibody used to validate KD mutants, p-SHP2 (Tyr580), p-GAB1 (Tyr642), p-MEK1/2 (Ser217/221), and p-ERK1/2 (Thr202/Tyr204) used to validate MAPK pathway activity, p-STAT3 (Tyr705), and p-STAT1 (Tyr701) used to validate STAT activation. **(D)** Western blot analysis of RNAi effect in PPP1CB::ALK models. GAPDH: loading control, p-ALK (Tyr1507) antibody used to validate retained ALK activity, p-SHP2 (Tyr580) and p-GAB1 (Tyr642), used to validate *shPTPN11*, p-STAT3 (Tyr705), used to validate *shSTAT3*. **(E)** Kaplan-Meier survival curves showing tumor induced mortality upon orthotopic intracranial injection of shRNA inhibited PPP1CB::ALK cells in NSG mice, groups are represented by individual curves, with a n=8 mice per group,; grey: PPP1CB::ALK^*shCtrl*^, dark petrol: PPP1CB::ALK^*shSTAT3*^, light petrol: PPP1CB::ALK^*shPTPN11*^; statistical significance determined by log-rank test, **: p-value<0.01, *:p-value<0.05.

SHP2 is best known as a regulator of MAPK signaling. To put these findings in a cellular context, we assessed activity levels of members of the SHP2/MAPK signaling axis by Western blot analyses of whole cell lysates. ALK-fusion expressing cells showed increased levels of GAB1, and to a lesser extent of MEK1/2 phosphorylation, whereas basal SHP2 phosphorylation levels appeared unaltered. Interestingly, increased p-SHP2 and p-GAB1 levels were observed in CLIP1::ROS1 and GOPC::ROS1 expressing cells, while downstream MAPK proteins remained largely unaltered. Notably, KIF21A::ROS1 cells revealed increased levels of total ERK1/2 protein (Figure 5C, Figure S4B).

In line with TF activity analyses (Figure 3F), elevated levels of phosphorylated STAT1 and STAT3 were detected in most ALK- and ROS1-fusion cells, except PPP1CB::ALK. KIF21A::ROS1 additionally showed increased total STAT1 levels, also detected as direct interactor of this fusion protein in the AP-MS/MS assay (Figure 5A,C, Figure S4B). *In vitro* kinase assays validated STAT3 as a direct substrate of ALK- and ROS1-fusions (Figure S4A) and subcellular fractionation Western-blots confirmed nuclear translocation of p-STAT3 (Figure S4C).

Whereas phospho-MS analysis did not reveal any active JAK kinases, the canonical kinases upstream of STAT1/3, the RTK adaptor proteins SHC1 for ALK-fusions (16) and SHC1/GRB2 for ROS1-fusions (17), both found in AP-MS/MS, have been reported to regulate STAT1/3, suggesting a non-canonical STAT activation (Figure 5A).

To analyze the effects of ROS1-fusions on SHP2 and STAT signaling acutely and independently of dominant-negative effects of the KD controls, short-term pharmacological inhibition (Entrectinib) was applied. After 4 hours of treatment, p-ROS1 was no longer detected, p-STAT1 and p-STAT3 levels were markedly reduced, p-SHP2 was ablated, and p-GAB1 levels were reduced (Figure S4D).

To further corroborate the direct link between increased p-SHP2 and p-STAT3 levels and oncogenic potential, we genetically suppressed *PTPN11* and *STAT3* in PPP1CB::ALK cells, deploying shRNA (Figure 5D). Intracranial injection of these cells into NSG mice resulted in a significantly increased survival of these mice compared to controls (Figure 5E), supporting our biochemical findings of ALK- and ROS1-fusions signaling via SHP2 and STAT3.

In summary, distinct biochemical and functional approaches confirmed SHP2 and STAT3 activation, underlying ALK- and ROS1-fusion tumorigenesis.

### Multiplexed protein mapping reveals spatial organization of Erk1/2 and Stat3 signaling in ALK- and ROS1-driven glioma murine models

To validate our findings in an independent *in vivo* model, we analyzed primary fusion-kinase-driven murine brain tumors generated by *in utero* electroporation (IUE) (6). This approach mimics pediatric cancer development more closely, with autochthonous *in vivo* tumor growth. To reveal the tissue- and subcellular distributions of ALK/ROS1 and downstream signaling effectors, we applied iterative indirect immunofluorescence imaging (4i) (18) on formalin-fixed, paraffin-embedded (FFPE) sections of tumor-bearing mouse brains (Figure 6A, S5A), as well as an age-matched healthy brain tissue. The tested samples, which in total comprised 320000 cells post QC, consisted of tumor areas (ALK or ROS1 positive) and non-tumor brain tissue, allowing for intrasample comparison. The expression of ALK or ROS1, p-Shp2, p-Erk1/2, and p-Stat3 at the single cell level was elevated in ALK/ROS1-positive areas compared to the negative neighboring areas (Figure 6B, Figure S5B). Increased p-Stat3 levels were detected within the nuclear area of ALK-or ROS1-fusion positive cells (Figure 6A), indicative of nuclear translocation and a functional upregulation of Stat3 activity.

**Figure 6:**
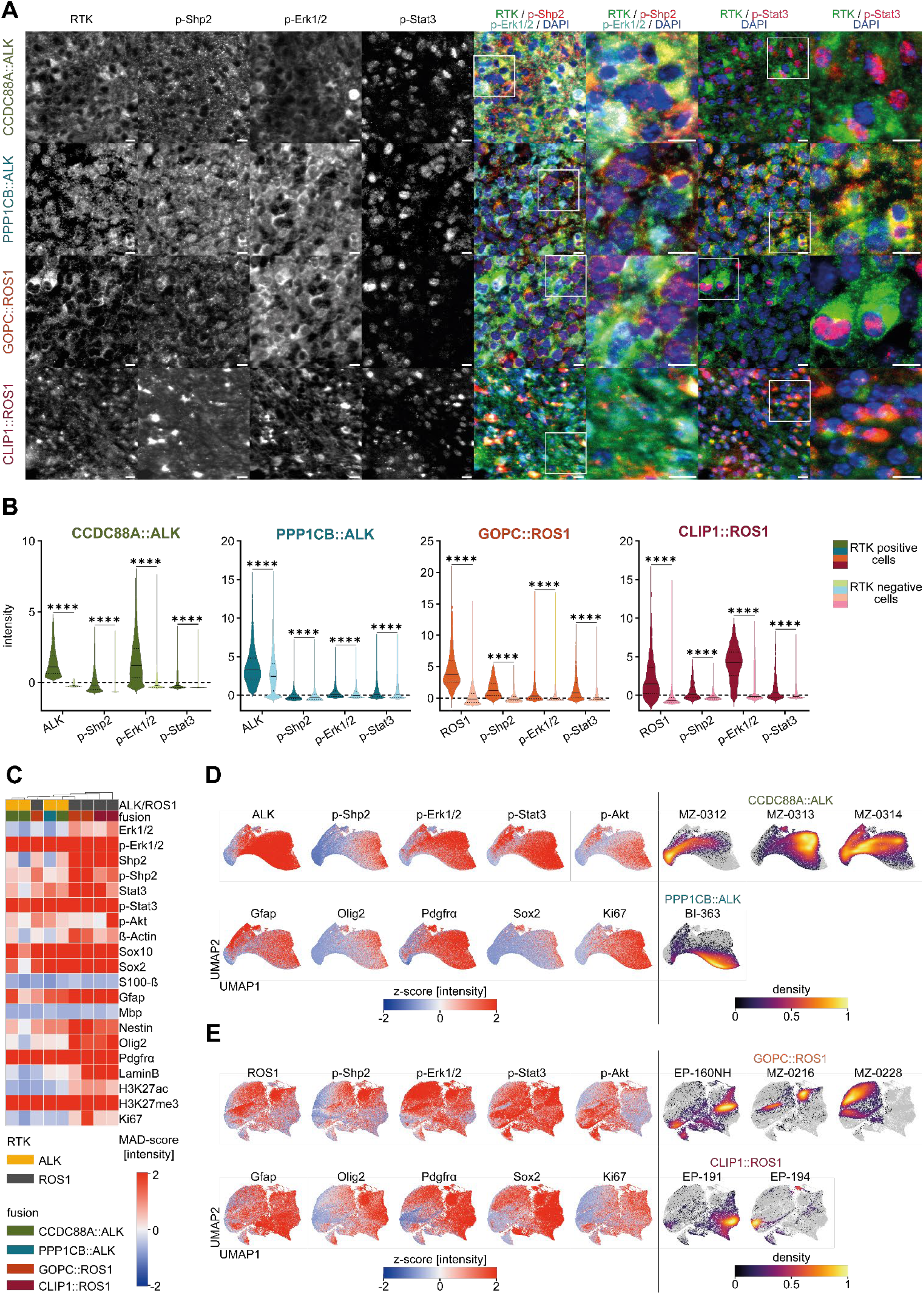
Spatial profiling reveals upregulated Stat3 and Shp2/Mapk signaling in primary ALK- and ROS1 murine tumors. **(A)** Illustrative region of analyzed brain tumors highlighting signaling marker expression; scalebar: 10µm. **(B)** Single cell intensity measurements for indicated antibodies detected by multiplex IF; x-axis: probed proteins, y-axis: intensities; visualization: violin plot, indicating interquartile range and median, Kruskal-Wallis-test, post-hoc Dunn-Bonferroni test: ****:p-value<0.0001. **(C)** Unsupervised hierarchical clustering of analyzed tumors based on MAD normalized average marker intensities, column-wise dendrogram clustering based on Euclidian distance and complete linkage; color gradient: MAD-score high (red) to low (blue). **(D-E)** UMAP representation of marker intensities at single cell resolution in **(D)** ALK-fusions and **(E)** ROS1-fusions, normalized to healthy control; left: UMAPs indicating marker intensities, right: density plots indicating localization of fusion samples within multidimensional space; color gradient: normalized z-score high (red) to low (blue); inferno color gradient: fusion distribution within UMAP space.

Fluorescence intensity levels of the signaling proteins examined varied between different fusions, corroborating fusion-specific pathway activities. Unsupervised hierarchical clustering of normalized mean marker expression revealed clustering driven by the RTK, with ROS1-fusions showing higher p-Shp2 intensities compared to ALK-fusions. Furthermore, whereas ALK- and ROS1-fusions showed increased expression of neural precursor markers Sox2, Sox10, and PDGFRα, ROS1-fusion cells specifically displayed higher levels of the progenitor marker Nestin, the oligodendrocyte precursor marker Olig2, and the active histone mark H3K27ac. Detection of neural progenitor markers suggest that, albeit differences, ALK- and ROS1-expressing tumor cells resemble an early neurodevelopmental-like cell state (Figure 6C).

UMAP projection of marker intensities at the single-cell level revealed partially overlapping multidimensional expression patterns between ALK or ROS1 and p-Shp2, p-Erk1/2, and p-Stat3. Closer analyses highlighted the heterogeneity between the studied fusions, which is exemplified by the more pronounced overlapping regions in the UMAP space of p-Erk1/2 expression with CCDC88A::ALK cells compared to PPP1CB::ALK cells (Figure 6D). Furthermore, staining for neural lineage marker revealed comparable distribution in the multidimensional space between neural stem cell (Sox2 and Nestin) and oligodendrocyte precursor cell lineage marker (Sox10, Pdgfrα, and Olig2) and ALK or ROS1 expressing cells, whereas marker for astrocytes (S100β) or oligodendrocytes (Mbp) are underrepresented (Figure 6D,E, S5C,D). These projections further demonstrate the differential upregulation of Mapk and Stat3 signaling in ALK-or ROS1-fusion. Additionally, assessment of neural lineage markers revealed aberrant activation of early neurodevelopmental programs in ALK- and ROS1-expressing cells.

Western blot analysis of murine tumor cells confirmed upregulation of Mapk and Stat3 signaling compared to healthy control brain (Figure S6A), and short-term treatment of CLIP1:ROS1 cells with Entrectinib resulted in reduced p-Mapk and p-Stat3 levels (Figure S6B). Furthermore, Entrectinib or Stattic treatment in CCDC88A::ALK and CLIP1::ROS1 murine tumor-derived cells resulted in dose-dependent cell death, corroborating our findings of RTK and STAT3 being critical for ALK- and ROS1-fusion driven tumors (Figure S6C).

Further, RNA sequencing in Entrectinib-treated vs. untreated murine tumor-derived cells, validated the upregulation of motility gene expression programs in CCDC88A::ALK expressing cells (Figure S6D). Furthermore, CCDC88A::ALK and GOPC::ROS1 cells repressed neural developmental pathways, suggestive of phenotypic plasticity of these tumors, and in line with the spatial analyses with enrichment in early neural development markers (Figure S6E).

Combining IUE technology with 4i spatial profiling, we confirmed increased Shp2 and Erk1/2 activity – more pronounced in ROS1-fused tumors – and increased Stat3 activity in all ALK- and ROS1-fused tumors with single-cell resolution. Importantly, we confirmed upregulation of gene expression programs promoting locomotion in CCDC88A::ALK. Finally, advanced tissue analyses hint at a dysregulation of neural development, marked by the expression of early neural markers in ALK- and ROS1 fusion-positive murine tumors.

## DISCUSSION

The detection of RTK fusions in pediatric gliomas, in particular ALK, ROS1, MET and NTRK fusions in young children with IHG, provides a strong rationale for the upfront use of targeted therapies (5,6,9). Detected exclusively in tumor cells and involving an activated kinase, RTK fusions are ideal targets for therapeutic intervention. Several early clinical studies are underway, and many more have been conducted over the last decade on other cancers with similar alterations (19-22). However, it remains unclear which pediatric glioma patients will benefit the most from specific inhibitor treatments and for how long. Similar RTK fusions are detected in several pediatric and adult cancers, suggesting functional overlap, and potential fusion-, tissue and/or age-specific oncogenic effects. Therefore, it is critical to define the roles of these fusions in different tumor contexts, to determine whether, and how, their heterogeneity impacts tumor formation.

Through the review of the OpenPedCan Project data, we confirmed that ALK and ROS1 fusions are overall rare events enriched in, but not limited to, IHGs, thus supporting the screening of these fusions across all pediatric gliomas. Unlike MET fusions — often co-detected with other known and tumor defining glioma drivers, such as H3K27M alterations (9,12) — most tumors with ALK and ROS1 fusions lack additional relevant oncogenic drivers.

ALK and ROS1 are RTKs similar in their sequence and structure, whereas ALK and ROS1 oncoproteins are very heterogeneous, with >50 fusion partners described (reviewed in 23,reviewed in 24). Nevertheless, ALK and ROS1 fusions are associated with a broad, yet largely overlapping spectrum of tumors, and thus presumed to be biologically similar. Despite the diversity in fusion partners, some typical characteristics of ALK and ROS1 fusions have been identified in this and in previous studies. With the loss of the 5’ end of the RTK gene, the expression of the fusion protein is generally controlled by the promoter of the fusion partner. Further, the fusion partner often dictates the intracellular localization (Figure 7) and impacts the stability of the fusion proteins (25). Several fusion partners possess a coiled coil domain that promotes oligomerization (e.g. CCDC88A and GOPC), whereas others promote the formation of cytoplasmic protein granules (e.g.: EML4 (26)), either one critical to the fusion’s activity (reviewed in 27).

**Figure 7:**
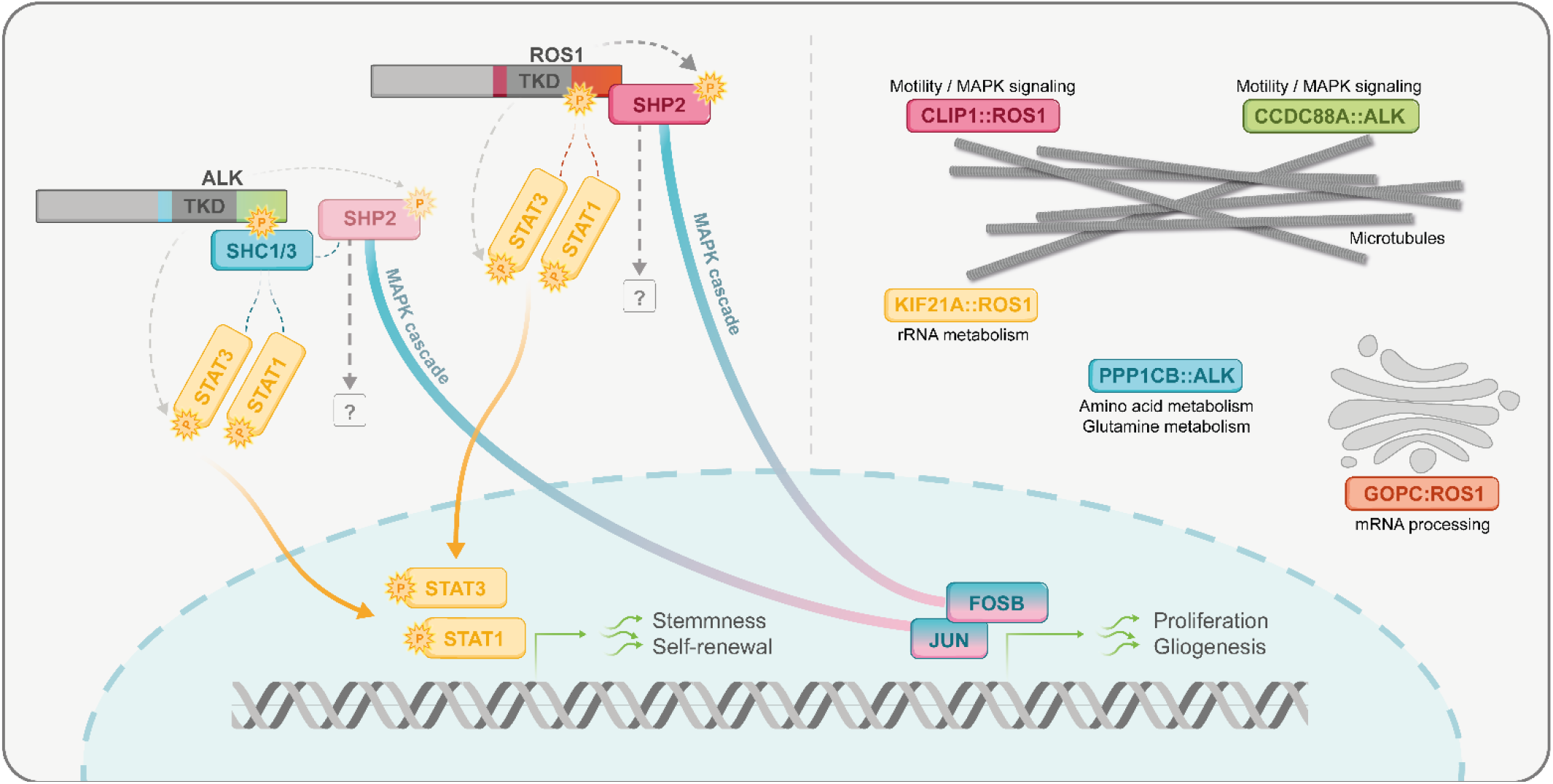
ALK- and ROS1-fusions with distinct cellular functions converge on STAT3 and SHP2 activation. Schematic representation of ALK- and ROS1-fusion driven signaling activation and subcellular localization. Left side: signaling pathways activated by ALK- and ROS1-fusions, dashed lines: Putative interaction based on published reports; dashed arrows: phosphorylation of substrates validated by *in vitro* kinase assays, TKD: tyrosine kinase domain; right side: subcellular localization of studied fusions, GO:terms specifically active in individual fusions annotated.

Using multi-omics profiling, we uncovered both convergent and fusion-specific functions, dictated in part by the fusion partner. Integration of transcriptomic and (phospho-)proteomic data generated on two ALK- and three ROS1-fusions revealed a heterogeneous landscape of differentially regulated genes and proteins. We agnostically compared the different fusions (and KD counterparts) and found that they separated into distinct subgroups, primarily, based on the presence of an ALK-or ROS1-fusion. Even accounting for potential experimental biases and use of different vector backbones, the functional heterogeneity among ALK- and ROS1-fusions was further confirmed in downstream analyses. The impact of the fusion partners was also evident, e.g. with the Golgi-associated GOPC::ROS1 fusion found to be associated with mRNA processing and metabolism, and the microtubule-interacting CLIP1::ROS1 and CCDC88A::ALK expressing cells revealing an upregulation of programs associated with cytoskeletal organization and cellular motility. Notably, we could prove that the increased motility caused by CLIP1::ROS1 was partly dependent on the MT domain of CLIP1, highlighting the relevance of the binding partners in fusion-specific cellular functions.

We identified SHP2 as overactivated and a direct interactor with all ROS1 fusions studied. ALK fusions were found in direct interaction with the adaptor proteins SHC1/3. Several other reports described an activation of SHP2 downstream ROS1 (28) and ALK fusions (29). SHP2 functions as a signaling integrator downstream of many RTKs, responsible in part for inducing and maintaining MAPK pathway activity. Biochemical and spatial analyses indicated stronger Ras/MAPK basal activation in ROS1 fusion-expressing cells. However, these changes were not as pronounced as the increase in SHP2 phosphorylation. In line with previous reports (28), our data suggest that SHP2 may also function via MAPK-independent non-canonical pathways. Conversely, SHP2 activation leading to MAPK activation has been reported to arise upon treatment with ALK inhibitors in NSCLC (30). Other studies point to the role of MAPK/ERK signaling in resistance to targeted NTRK inhibition in NTRK-fused tumors (31,32), and co-targeting RTK/SHP2 approaches have been proposed as a strategy to overcome MAPK reactivation and resistance to RTK targeted therapies (reviewed in 33).

The role of MAPK signaling downstream ALK- and ROS1-fusions should be further investigated, as its regulation appears to be manifold and potentially distinct between different fusions. MOFA analyses revealed upregulation of MAPK-associated TFs in most fusion models, whereas at a protein level, phosphorylation of MAPK effectors was detected primarily in CCDC88A:ALK and CLIP1::ROS1. Further, in murine tumors, we detected an increase in cytoplasmic (rather than nuclear) p-Erk, hinting at oncogenic cytosolic Erk signaling. For example, Erk has been shown to enhance activation of Stat3 within the cytosol by phosphorylation of Ser727 (34). We noted increased activation of STAT signaling, in particular STAT1 and STAT3, downstream all fusions, and confirmed p-STAT3 nuclear localization in the tissue context. Mechanistically, our data point towards direct phosphorylation of STAT3 by the fusion kinases, and the phospho-MS results support such a model of non-canonical, JAK-independent activation of STAT signaling (Figure 7).

We further deployed 4i and spatially confirmed an upregulation of STAT signaling and differential fusion-dependent SHP2/MAPK activation in murine tumor tissues. The spectrum of fusions detected in patients with IHG is quite heterogenous, with several patients presenting with unique fusions. Considering that individual fusions activate cellular signaling hubs to different extents, it is conceivable that similar advanced tissue-based analyses could be used in the future to provide insights on pathway activity at single-cell resolution. Further work should continue to dissect the biological interplay between fusion proteins and signaling networks, to understand how protein activity and gene regulatory programs might impact response to therapy.

In conclusion, in this study we provide detailed mechanistic information, working towards a better understanding of IHG biology. Our findings show that ALK and ROS1 fusions found in IHGs convergence in downstream STAT and MAPK pathway activity. The knowledge gained can be used to refine diagnostic approaches and develop new treatment strategies for children with IHGs.

## Acknowledgements

We thank Antje Dittman and Jonas Grossmann from the Functional Genomic Center Zurich for their support with the establishment and optimization of phospho-proteomics analyses. We further thank Arpan Rai and Adrian Tschan from Apricot Therapeutics AG for their technical support. This work was possible due to grant support from: Swiss National Science Foundation (#PCEFP3_194620); Heidi Ras Stiftung; Swiss Bridge Award; Fonds zur Förderung des akademischen Nachwuchs, Universität Zürich; Promedica Stiftung.

## Author contributions

A.S.G.S., A.P., A.S.B. designed and conceived the study with critical input from all authors. Most experiments were conducted by A.P. and A.S.B., with support from: N.H. and S.C. for *in vivo* work; R.D., E.B.C., L.B.G. and L.B. for *in vitro* validation experiments; S.Y., B.C. and M.B. for cell motility experiments. IUE *in vivo* models were performed by N.H. under supervision of M.Z. Multi-omic data integration/MOFA analyses performed by J.T.D., under supervision of O.A.. RNAseq data analyses done additionally by C.M. and A.J.D.M.. The first draft of the manuscript was written by A.S.G.S., A.P. and A.S.B., with subsequent edits from the other authors.

## Data availability

The mass spectrometry proteomics data have been deposited to the ProteomeXchange Consortium via the PRIDE partner repository with the dataset identifier PXD059984.

The transcriptomic data have been deposited to the Zenodo repository with the digital object identifier 10.5281/zenodo.14848911.

## METHODS

### Cellular models

Immortalized normal human astrocytes (TERT/E6/E7) (iNHA), iNHAs stably expressing flag-tagged CCDC88A::ALK, or PPP1CB::ALK, and EV were a gift from Dr. Hawkins (5). ALK-fusion controls and ROS1-fusion expressing stable lines were generated by lentiviral transduction as described previously (5) and mCherry-positive cell sorting or Puromycin selection [10µg/ml], respectively. iNHA were adherently cultured on TC-treated plastic ware in DMEM (Thermo Fisher #119600044) supplemented with 10% FBS (Sigma Aldrich #S0615-500ML), 1mM GlutaMAX (Thermo Fisher # 35050061), 1mM sodium pyruvate (Thermo Fisher #11360039), 50U/ml Penicillin-Streptomycin (Thermo Fisher #15070063) and passaged using Trypsin-EDTA (BioConcept #5-51F00-I). Murine tumor derived cells were cultured in spheres in low attachment flasks (Thermo Fisher #174951, #174952) or multi-well plates (Greiner BIO-ONE #657970, #662970) in a 1:1 mix of Neurobasal-A (Thermo Fisher #10888022) and DMEM/F-12 (Thermo Fisher #11330032) supplemented with 10mM HEPES buffer solution (Thermo Fisher # 15630056), 1mM sodium pyruvate (Thermo Fisher #11360039), 0.1M MEM non-essential amino acids (Thermo Fisher # 11140050), 1mM GlutaMAX (Thermo Fisher # 35050061), 50U/ml Penicillin-Streptomycin (Thermo Fisher #15070063), 2% B-27 without vitamin A (Thermo Fisher #12587010), 2µg/ml heparin (Stemcell Technologies #07980), 10ng/ml PDGF-AA (Peprotech #100-13A), 20ng/ml recombinant human bFGF (Peprotech #100-18B), and 20ng/ml recombinant human EGF (Peprotech #AF-100-15) (6). Spheres were separated using Accutase (Stemcell Technologies #07922). Maintenance and splitting of all cell lines was done at 37°C and 5%CO_2_. Cell line identity was validated by short tandem repeat analysis and cells were tested negative for mycoplasma biannually.

### *In vivo* experiments

For orthotopic intracranial injections 6 – 8 weeks old male and female NOD.Cg-*Prkdcscid Il2rgtm1Wjl*/SzJ (NSG) mice (Charles River #005557) were used, and 8 animals were randomly assigned to either a control or experimental group. Mice assigned to either group were housed together, no mixing of control and experimental mice was performed. Mice were fixed in a stereotactic frame, and 200,000 ALK-or ROS1-fusion cells were injected in the brain cerebral cortex. Animals were monitored 3 to 5 times a week and euthanized, via CO_2_ asphyxiation as per license, in the presence of physiological signs of a brain tumor (weight loss, domed head, hunched posture, scruffy appearance) or one year post injection in the control group. Single female mice were combined with other females from the same group to prevent single housing. Male mice were single housed if cage mates reached endpoint. Post euthanasia, brains were collected and fixed using 4% formaldehyde, pH 7, (ROTI Histofix, Carl Roth, Cat.NR: P087.5) for immunohistochemistry. All procedures were performed under License ZH108/2020 approved by the Zurich Cantonal Animal Experimentation Commission. *In utero* electroporation was performed as previously described (6). Briefly, CD1 embryos (strain: RjOrl:SWISS) were injected with previously described plasmids together with a guide RNA targeting *Tp53* into the lateral ventricle and electroporated *in utero* at E14.5. Three days post birth, successful integration was validated using bioluminescence imaging on an IVIS imager (PerkinElmer). Mice were sacrificed according to human endpoint criteria. Brains were extracted at necropsy, sagittally cut through the tumor and one part fixed using 4% formaldehyde, pH 7, (ROTI Histofix, Carl Roth, Cat.NR: P087.5) The other tumor half was extracted, mechanically dissociated, strained through a 40-µm cell strainer and plated in the above described medium to generate cell lines (6).

### Multi-Omics Factor Analysis (MOFA)

Unsupervised integration of the RNA, proteome, phosphoproteome (RNA/prot/phospho) layers was performed by Multi-Omics Factor Analysis (MOFA2 package) as previously described (15). MOFA decomposes multi-omics datasets into hidden molecular patterns, so called latent factors, driving variance across individual molecular modalities (RNA/prot/phospho) and cell lines, respectively. Mathematically, each factor ordinates the samples along a one-dimensional axis centered at zero. Scored ranking of genes (weighted by their contribution), proteins and PTMs is extracted for each factor and data modality which is used for subsequent downstream analyses. Features with no association to a given factor have MOFA scores close to zero, while features with strong associations have large absolute MOFA scores. MOFA was applied separately to ALK and ROS1 fusion cell type samples. Upon filtering out the data for missing values and features without variance across cell lines (ANOVA, p-value cut-off = 0.01), 9177 or 8548 transcripts, 4248 or 5554 proteins, and 8164 or 7799 phosphorylated proteins for ALK-or ROS1-fusions respectively, were used to perform MOFA (Figure 3B,C).

### Biological activity inference (DecoupleR)

Transcription factor activities were inferred using the DecoupleR tool (35). DecoupleR contains different statistical methods to extract biological signatures from prior knowledge within a unified framework. For this study, a Univariate Linear Model (ULM) was applied. Transcription factor (TF) activity was computed from RNAseq data, using RNA MOFA scores (Factor1 in ALK and Factors 3 and 5 in ROS1 samples) as input, and TF-target collection from CollecTRI database which compiles signed TF – target gene interactions from 12 different resources (36). Only TFs with at least 10 measured targets were included.

### Cell Motility Assay

iNHAs were seeded at 7,000 cells/200µl per well in treated (ibiTreat) 8-well chamber microscopy slides (ibidi #80862). Cells were cultured in the above described iNHA medium for 24 hours. Cell nuclei were labeled using the CellLight Nucleus-GFP, BacMam 2.0 (Thermo Fisher #C10602) according to the manufacturer’s protocol, with overnight incubation for optimal expression. Live-cell imaging was conducted on a Nikon ECLIPSE Ti2 inverted widefield microscope, capturing images every 6 minutes for 12 hours at 10× magnification. Image analysis for motility and tracking was performed using ImageJ software (37).

### Imaging

Live cell imaging was conducted to track nuclear fluorescence using a Nikon Ti2 inverted microscope, equipped with a 20X/0.4 NA air objective and fluorescence filter sets for GFP and Cy3. Cells were imaged over a 12-hour period, with frames captured every 3 minutes (20 frames per hour). To mitigate phototoxicity and photobleaching, excitation light was applied only during camera acquisition, with both excitation intensity and exposure time carefully calibrated to maximize the signal-to-noise ratio while minimizing these effects. No detectable bleaching was observed over the 12-hour recording, indicating that phototoxicity effects were minimal.

### Movie Preprocessing

Before tracking, movies were preprocessed to reduce file size and improve the signal-to-noise ratio and contrast, essential for more robust object identification during the tracking step. First, movies were downscaled by a factor of 4 through pixel binning (local averaging). Background fluorescence from the culture medium was then removed on a frame-by-frame basis using median filtering: a median-filtered version of each frame, created with a 25-pixel kernel size, was subtracted from the original image. This approach effectively functions as a high-pass filter. Second, contrast was enhanced by limiting and redistributing the dynamic range to 8-bit. Specifically, the global background intensity was estimated as the 50th percentile of the entire movie’s intensity—a reasonable approximation for sparse fluorescence signals. This background value was used to limit the data range: after normalizing to the background value, pixel intensities were constrained to a range of 1–5, effectively setting a minimum limit at the background level and a maximum limit at 5 times this background value. Finally, values were normalized to a 0–255 range and converted to 8-bit.

### Tracking

Nuclei tracking was performed by scripting the Fiji plugin TrackMate (38), available at *ImageJ Wiki* https://imagej.net/plugins/trackmate/, using the tracking algorithm developed by Jaqaman *et al. (39)*. This algorithm consists of three main steps: object detection, step linking, and trajectory stitching. In the object detection step, nuclei were identified in each frame using a Laplacian of Gaussian (LOG) algorithm, which localizes each nucleus at its center point (refer to plugin documentation for details). The parameters used for detection were a radius of 11μm (estimated object size) and a threshold of 1, corresponding to the minimum intensity required for object detection. For step linking and trajectory stitching, the plugin’s SparseLAPTrackerFactory() was applied, utilizing the LAP (Linear Assignment Problem) algorithm (see plugin documentation). The following parameters were set for tracking:LINKING_MAX_DISTANCE (maximum step size) at 30μm, GAP_CLOSING set to ‘False’, ALLOW_TRACK_SPLITTING set to ‘False’, and ALLOW_TRACK_MERGING set to ‘False’. These settings effectively disable the ability to stitch together spatially separated trace fragments, allowing us to retain only traces of objects that are robustly detected in each frame. Additionally, in splitting events, traces of mother and daughter cells are treated as distinct objects, ensuring separate tracking for each.

### Data Plotting and Post-Processing

For downstream data analysis and visualization, we considered only traces with a minimum duration of 60 minutes and non-zero displacement (i.e., non-stacked objects). Further details on the plotting parameters can be found in the figure caption.

### Indirect iterative immuno fluorescence

Staining and analyses, including UMAP projections, of FFPE embedded brain slices were performed by Apricot Therapeutics according to Kramer *et al*. (40).

### Plasmids

Flag-tagged CCDC88A::ALK, PPP1CB::ALK, and EV pLVX plasmids were a gift from Dr. Hawkins (5). HA-StreptagII-tagged GOPC::ROS1, CLIP1::ROS1, and KIF21A::ROS1, or HA-StreptagII only sequences were cloned into pLVX-IRES-mCherry by Genscript USA to generate iNHAs expressing ROS-fusion transgenes. mCherry sequence was replaced by Puromycin resistance from pSpCas9(BB)-2A-Puro (PX459) (Addgene #48139 (41)) by seamless cloning (In-Fusion Snap Assembly, Takara # 638945) using the following primer: mCherry removal forward: 5’-TGAACGCGTCTGGAACAATCAAC-3’, mCherry removal reverse: 5’-ATTATCATCGTGTTTTTCAAAGGAAAACCAC-3’; Puromycin insert forward: 5’-AACACGATGATAATACCCGCCATGGAGATCGAGTG-3’, Puromycin insert reverse: 5’-TGTTCCAGACGCGTTGGCGAAGGCGATGGGGGTC-3’. Mutant controls were generated using directional mutagenesis (In-Fusion Snap Assembly, Takara # 638945) using the following primer: ALK-fusion kinase dead forward: 5’-GGCTGTGATGACGCTGCCTGAAGTGTGC-3’, ALK-fusion kinase dead reverse: 5’-ACGGTCATCACAGCCACTTGCAGGGG-3’; ROS1-fusion kinase dead forward: 5’-GGCCGTGATGACCCTGAAGAAGGGCAGC-3’, ROS1-fusion kinase dead reverse: 5’-AGGGTCATCACGGCCACCTTGATCTCG-3’; CLIP^ΔTUB^ forward: 5’-TAGGCGAAGAGGATGGTTCGGTGGCAGGA-3’, CLIP^ΔTUB^ reverse: 5’-CATCCTCTTCGCCTATGGGTTCATCTAAAACA-3’; sequencing primer to validate correct mutations: CMV-forward: 5’-CGCAAATGGGCGGTAGGCGTG-3’, IRES-reverse: 5’-TATAGACAAACGCACACCG-3’. CCDC88A::ALK, GOPC::ROS1, GOPC::ROS1^KD^, CLIP1::ROS1, and ROS1^ex35^ were cloned from pLVX vectors into pT2K-CAGGS-IRES-Luc (6) by seamless cloning (In-Fusion Snap Assembly, Takara # 638945) using ALK-IUE-forward: 5’-TTTTGGCAAAGAATTGACTACAAAGACGATGACGACAAGGAG-3’, ALK-IUE-reverse: 5’-ATTGATCCCGCTCGAGATCAGGGCCCAG-3’, GOPC:ROS1-IUE-forward: 5’-TTTTGGCAAAGAATTACCATGTCGGCGGGCGGTCC-3’, GOPC::ROS1-IUE-reverse: 5’-ATTGATCCCGCTCGATCAAGCGTAATCTGGAACATCGTATGGGTAATCAGACCCATCTCCATATCCACT-3’, CLIP1::ROS1-IUE-forward: 5’-TTTTGGCAAAGAATTACCATGAGTATGCTAAAGCCAAGTGGGC-3’, CLIP1::ROS1-IUE-reverse: 5’-ATTGATCCCGCTCGACAAGCGTAATCTGGAACATCGTATGGGTTAATCAGACCCATCTCCATATCCACT-3’, ROS1^ex35^-IUE-forward: 5’-TTTTGGCAAAGAATTTGGCATAGAAGATTAAAGAATCAAAAAAGTGCCA-3’, ROS1^ex35^-IUE-reverse: 5’-ATTGATCCCGCTCGATTAATCAGACCCATCTCCATATCCACT-3’. PT2K-IRES-LUC was digested using *EcoRI* (NEB # R3101S) and *XhoI* (NEB # R0146S). Sequencing primer to validate correct insertions: PT2k-forward: 5’-GCTCTAGCTAGAGCCTCTGCTAACC-3’ and PT2K-reverse: 5’-CCAAAAGACGGCAATATGGTG-3’. shRNAs targeting either *PTPN11* (Sigma Aldrich Mission shRNA; TRCN0000355818), *STAT3* (Sigma Aldrich Mission shRNA; TRCN0000329887), or a scrambled control (Addgene #162011) were transduced into PPP1CB::ALK cells using lentiviral approaches and selected using 250µg/ml Geneticin for shPTPN11 and shSTAT3, or 20µg/ml Puromycin for shCtrl.

### Histology and immunohistochemistry

4µm thick sections of FFPE embedded brains were cut and stained for H&E following manufacturer’s instructions (Sakura #4900) using an automated device (Sakura, Tissue-Tek Prisma). IHC stainings with the following antibodies and according to manufacturer’s instructions were performed on an automated device (Leica, BOND; Roche, Ventana): mouse-α-ALK (1:10, 30min; Leica Biosystems #NCL-L-ALK), rabbit-α-p-ERK1/2 (1:200, 30min; CST #4370), rabbit-α-p-SHP2 (1:100, 30min; Invitrogen #PA5-114642), rabbit-α-p-STAT3 (1:100, 44min; CST #9145), rabbit-α-KI67 (1:40, 60min; Abcam #ab16667), and rabbit-α-ROS1 (1:100, 32min, Ventana; CST #63452).

### RNA extraction

iNHA cells were grown for 24 without FBS and harvested by trypsinization. IUE cells were grown in suspension for 24 hours supplemented with either 100nM Entrectinib or DMSO before harvesting by centrifugation and RNA was extracted using the RNeasy Plus Mini Kit (Qiagen #74134) according to manufacturer’s protocol. RNA sequencing was conducted at the Functional Genomic Center Zürich (FGCZ).

### Library preparation

The quality of the isolated RNA was determined with a Fragment Analyzer (Agilent, Santa Clara, California, USA), and samples with a 260 nm/280 nm ratio between 1.8–2.1 and a 28S/18S ratio within 1.5–2 were further processed. The Illumina® Stranded mRNA Prep, Ligation kit (Illumina, Inc, California, USA) was used in the succeeding steps. Briefly, total RNA samples (100-1000 ng) were poly A enriched and then reverse-transcribed into double-stranded cDNA. cDNA samples were fragmented, end-repaired and adenylated before ligation of an anchor. Fragments containing the anchor on both ends were selectively enriched with PCR at the same time adding the index with UDI. The quality and quantity of the enriched libraries were validated using the Fragment Analyzer (Agilent, Santa Clara, California, USA). The product is a smear with an average fragment size of approximately 260 bp. The libraries were normalized to 10nM in Tris-Cl 10 mM, pH8.5 with 0.1% Tween 20.

### Cluster Generation and Sequencing

The NovaseqX (Illumina, Inc, California, USA) was used for cluster generation and sequencing according to standard protocol. Sequencing was paired end at 2 X150 bp. All relevant data have been deposited to the Zenodo repository with the digital object identifier 10.5281/zenodo.14848911.

### RNAseq preprocessing

RNA sequencing (RNA-seq) data from iNHA were analyzed using the bioinfo-pf-curie/RNA-seq pipeline (version 4.1.0) (42), with the human genome reference Hg19. For mouse RNA-seq data, analysis was performed using the nf-core/rnaseq pipeline (version 3.14.0; https://zenodo.org/records/10471647) (43), implemented in Nextflow. The reference genome utilized was GRCm39 (Mus musculus). RNA-seq data analysis followed the default parameters provided by the nf-core/rnaseq.

### RNA-seq data analysis

RNAseq matrix was filtered and normalized using standard Bioconductor DESeq2 package (44). Downstream analysis was only restricted to protein-coding genes and all genes having no count across all samples were filtered out before normalization. Raw counts were normalized using the rlog Transformation method from bioconductor DESeq2 package. The 1000 most variable genes across all ALK- and ROS1-fusion cell lines (lowest p-value from ANOVA analysis) were kept for exploratory analysis (PCA and hierarchical clustering analysis using FactoMineR (45) and ComplexHeatmap (46) R packages, respectively. Differential gene expression analyses were performed using DESeq function of the DESeq2 package using the filtered raw counts. The resulting p-values were corrected using the Benjamini and Hochberg method.

### Protein extraction

For mass spectrometry iNHA cells were grown for 24 without FBS and harvested by trypsinization. IUE cells were grown in suspension for 24 hours supplemented with either 100nM Entrectinib or DMSO before harvesting by centrifugation. Proteins were extracted using RIPA buffer (50mM Tris-HCl pH 7.4, 150mM NaCl, 1% IGEPAL CA-630 (NP-40), 0.5% sodium deoxycholate, and 0.1% SDS) supplemented with 1mM PMSF, 1mM Na_3_VO_4_, phosphatase (Roche # 4906845001) and protease (Roche # 11836170001) inhibitor tablets. Protein concentration was determined using a BCA assay (Thermo Fisher # 23225). For affinity purification mass spectrometry iNHA cells were grown 24 hours without FBS, harvested by trypsinization and proteins were extracted using 20mM Tris-HCl pH 7.5, 1% Tween-20, 0.3% IGEPAL-CA 630, supplemented with 1mM PMSF, 1mM Na_3_VO_4_, phosphatase (Roche # 4906845001) and protease (Roche # 11836170001) inhibitor tablets. Protein concentration was determined using a BCA assay and adjusted to equal concentrations (Thermo Fisher # 23225). Lysates were precleared using magnetic protein G beads (Thermo Fisher #10003D) for 1 hour at 4°C. Pulldowns were performed over night at 4°C using 20µl bead slurry/mg lysate magnetic anti-flag beads (Sigma Aldrich #M8823) for ALK-fusions or StreptactinXT beads (IBA # 2-5090-002) for ROS1-fusions, respectively. Following 2 washes using lysis buffer followed by 2 washes using 1xTBS (20mM Tris-HCl pH7.5, 150mM NaCl) bound proteins were eluted twice using 0.1mg/ml FLAG peptide (Sigma Aldrich #F3290) for ALK-fusions or 2.5mM desthiobiotin (IBA #2-1000-002) for ROS1-fusions respectively. Mass spectrometry was conducted at the Functional Genomic Center Zürich (FGCZ).

### LC-MS/MS sample preparation

Eluted proteins from AP-MS experiments were precipitated with trichloroacetic acid (TCA; Sigma-Aldrich) to a final concentration of 5% and washed twice with ice-cold acetone. Samples were resuspended in digestion buffer (10 mM Tris-HCl, pH 8.0, 2mM CaCl_2_), reduced with 5mM TCEP (tris(2-carboxyethyl)phosphine) and alkylated with 15mM chloroacetamide at 30°C for 30min, followed by overnight digestion with 500ng of Sequencing Grade Trypsin (Promega) at 37°C. Samples for phospho- and global proteome analyses were reduced and alkylated before SP3-based protein purification, digestion and peptide clean-up using a KingFisher Flex System (Thermo Fisher) and Carboxylate-Modified Magnetic Particles (GE Life Sciences; GE65152105050250, GE45152105050250) (47,48). Beads were conditioned following the manufacturer’s instructions, consisting of 3 washes with water at a concentration of 1µg/µl. For each sample, 60µg of protein (according to absorption measurement) were used and diluted with 100% ethanol to a final concentration of >50% ethanol. The following steps were carried out on the robot: collection of beads from the last wash, protein binding to beads, washing of beads in wash solutions 1-3 (80% ethanol) and peptide elution from the magnetic beads using MilliQ water. Protein digestion with Trypsin at a peptide-to-protease ratio of 50:1 was carried out on a Thermoshaker overnight at 37°C. The digest solution and water elution were combined and dried to completeness. Afterwards, 250µg TMT18pro reagent (Thermo Fisher #90110) was dissolved in 15μl of anhydrous acetonitrile (Sigma-Aldrich) and added to 60µg peptides in 45µl of 50mM TEAB, pH 8.5. The solution was gently mixed and incubated for 60min at room temperature. The reaction was quenched by adding 3.5µl of 5% hydroxylamine (Thermo Fisher). The combined TMT sample was created by mixing equal amounts of each TMT channel. Labeled peptides were fractionated offline using high pH reverse phase chromatography. In short, peptides were separated on an XBridge Peptide BEH C18 column (130Å, 3.5µm, 4.6mm X 250mm, Waters) using a 72min linear gradient from 5-40% acetonitrile/9 mM NH_4_HCO_2_. Every minute a new fraction was collected and concatenated to 12 final fractions and a small amount was taken for global proteome analysis before drying to completeness. Phosphopeptide enrichment was performed on the KingFisher Flex System (Thermo Fisher) (48) using PureCube Fe-NTA MagBeads (Cube Biotech). Beads were conditioned following the manufacturer’s instructions, consisting of 3 washes with 200µl of binding buffer (80% acetonitrile, 0.1% TFA). Each fraction was dissolved in 150µl binding buffer. Enrichment was carried out using the following steps: washing of the magnetic beads in binding buffer (5min), binding of the phosphopeptides to the beads (30min), washing the beads in wash 1-3 (binding buffer, 3 min each) and eluting peptides from the beads (50µl 2.5% NH4OH in 50% acetonitrile, 10min). Each elution was combined with 30µl neutralisation solution (75% acetonitrile, 10% formic acid) and samples were dried to completeness and re-solubilized in 10µl of 3% acetonitrile, 0.1% formic acid for MS analysis. Enrichment of phospho-tyrosine containing peptides was assessed as described previously (49) with some modifications. In short, 12μg 4G10 (Millipore #16-204) and 12μg pY-1000 (CST #8954) were conjugated to Protein G beads and incubated with TMT-labeled peptides in 100mM Tris– HCl, 1% IGEPAL-C630 rotating at 4°C overnight. After acid elution, the peptides were subjected to a secondary enrichment step using an Fe-NTA spin column (Thermo Fisher) according to the manufacturer’s protocol. Eluted peptides were dried to near-completeness and loaded onto Evotips, according to the manufacturer’s protocol.

### LC-MS/MS data acquisition

Mass spectrometry analyses of AP-MS, and global and phospho-proteome samples were performed on an Orbitrap Exploris 480 mass spectrometer (Thermo Fisher) equipped with a Nanospray Flex Ion Source (Thermo Fisher) and coupled to an M-Class UPLC (Waters). Solvent composition at the two channels was 0.1% formic acid for channel A and 0.1% formic acid, 99.9% acetonitrile for channel B. Column temperature was 50°C. Peptides were loaded on a commercial nanoEase MZ Symmetry C18 Trap Column (100Å, 5µm, 180µm x 20mm, Waters) connected to a nanoEase MZ C18 HSS T3 Column (100Å, 1.8µm, 75µm x 250mm, Waters). The flow rate was set to 300 nl/min. For global- and phospho-proteome analyses, TMT-labeled (phospho-)peptides were separated using a gradient from 5 to 22 % B in 80min (40min for unlabeled AP-MS samples) and 22 to 32% B in additional 10min (5min for AP-MS samples). The column was cleaned after each run by increasing to 95 % B and holding 95 % B for 10min prior to re-establishing loading condition for another 10min. pTyr-peptides were separated on an Evosep One system using the 15 SPD method and commercial PepSep C18 column (1.5µm, 150µm x 150mm). The mass spectrometer was operated in data-dependent mode (DDA) with a maximum cycle time of 3s, spray voltage set to 2.4kV, funnel RF level at 40%, heated capillary temperature at 275°C, and Advanced Peak Determination (APD) on. Full-scan MS spectra (350−1’500m/z) were acquired at a resolution of 120’000 at 200m/z after accumulation to a target value of 3000000 or for a maximum injection time of 45ms. Precursors with an intensity above 5’000 were selected for MS/MS. Ions were isolated using a quadrupole mass filter with 0.7m/z isolation window for TMT-labeled samples (1.2m/z for unlabeled AP-MS samples) and fragmented by higher-energy collisional dissociation (HCD) using a normalized collision energy of 32% (30% for unlabeled AP-MS samples). HCD spectra were acquired at a resolution of 60’000 or 45’000 (phospho, pTyr), or 30’000 with (global proteome TMT) or without (AP-MS) turboTMT on. Maximum injection time was set to 200ms (phospho), 119ms (AP-MS), 300ms (pTyr) or automatic (global proteome). The normalized automatic gain control (AGC) was set to 100%. Charge state screening was enabled such that singly, unassigned and charge states higher than six were rejected. Precursor masses previously selected for MS/MS measurement were excluded from further selection for 20s, and the exclusion window was set at 10ppm. The samples were acquired using internal lock mass calibration on m/z 371.1012 and 445.1200. The mass spectrometry proteomics data were handled using the local laboratory information management system (LIMS) (50) and all relevant data have been deposited to the ProteomeXchange Consortium via the PRIDE (http://www.ebi.ac.uk/pride) (51) partner repository with the data set identifier PXD059984.

### LC-MS/MS data analysis

MS data of TMT-labeled samples were processed for identification and quantification using Proteome Discoverer 2.5.0.400 (Thermo Fisher Scientific). Spectra were searched against Ensembl’s Homo sapiens Genome assembly: GRCh38 concatenated to its reversed decoyed fasta database using Sequest HT with FDR calculation done using Percolator. FDR thresholds were specified as 1% on psm and on peptide level. TMT modification on peptide N-termini and Lysine side chains as well as carbamidomethylation of cysteine were set as fixed modification, while methionine oxidation, and phospho (STY) were set as variable. Enzyme specificity was set to trypsin/P allowing a minimal peptide length of 6 amino acids and a maximum of two missed cleavages. Precursor tolerance was set to 10ppm and fragment ion tolerance was set to 20mmu. Reporter ion intensities were extracted with 20ppm integration tolerance. For peptide and protein quantification the co-isolation filter was set to 50 % and the average signal-to-noise threshold to 10. Site localization probability was scored using the ptmRS mode with a probability cut-off of 0.75. The peptideGroups.txt output of Proteome Discoverer was taken and further analyzed using the prolfqua R-package (52). In brief, peptides were accepted if they carried a phosphorylation and the reported abundances for all TMT channels were filtered for a minimum abundance of 1. Further, these reporter channel abundances were log_2_-transformed and normalized with a robust z-score transformation. Group comparisons (contrasts) were evaluated with a moderated Wald-test with pooled variance (as implemented in the limma R-package (53)). The resulting p-values were adjusted for multiple testing using BH-method. Global proteome results were analyzed in the same way, but the input file was the “Proteins.txt” file from Proteome Discoverer. Here we filtered for “IsMasterProtein” as well as for FDR Confidence combined equals “High”. AP-MS samples were processed for identification and quantification using, MSFragger (version 3.3) (54) and Philosopher (version 4.4.0) (55). Spectra were searched against a Uniprot Homo sapiens reference proteome (UP000005640), concatenated to its reversed decoyed fasta database and common protein contaminants. For the closed search settings, strict Trypsin digestion with a maximum of 2 missed cleavages was set. Carbamidomethylation of cysteine was set as fixed modification, while methionine oxidation and N-terminal protein acetylation were set as variable. Label free quantification and match between run option were enabled. Protein abundance estimates were extracted from the intensity columns of the combined_protein.tsv file and processed for scoring potential interactions between observed proteins and the bait protein using SAINTexpress software (56). Data conversion into SAINTexpress compatible format, and visualizations were performed using the R package prolfqua (52).

### Phospho and Proteome data analysis

For exploratory analysis for proteome and phospho data the 1000 most variable proteins and PTMs across all ALK and ROS1 cell lines were subjected to PCA and hierarchical clustering analysis using FactoMineR (45) and ComplexHeatmap R packages (46), respectively. The differential protein and PTM expression analysis was carried out by t-test function (from rstatix R package (57)) adapted to data with missing values. Finally, to identify unique proteins or PTMs specificity score was computed for all proteins and PTMs. It is defined as the ratio between the percentage of replica in which a given protein/PTM is detected in one cell lines and the total percentage in all cell lines: a score of 1 means that the protein/PTMs is only detected in one cell line while a score of 0 means that the protein/PTM is absent. PTPN11/SHP2 Y584 annotation had a low site-localization probability, which was later experimentally confirmed to actually be Y580.

### Drug titrations

2500 cells were seeded in round bottom ultra-low attachment 96well plates (Sigma Aldrich #CLS4515-5EA). 24 hours later 1 volume of medium containing double the indicated drug concentration, or concentration adjusted DMSO, was added and the cells were incubated for 72h at 37°C with 5%CO_2_. Drugs used were entrectinib (Selleckchem #S7998), trametinib (Selleckchem #S2673), and stattic (Selleckchem #S7024). Viability readout was performed using a metabolic assay (CellTiter-Glo® 3D, Promega #G9681) according to manufacturer’s instructions. Recorded luminescence was normalized to DMSO samples and set to 100%. 3 independent biological replicates with 3 technical replicates each were performed.

### I*n vitro* kinase assay

iNHAs were grown in the absence of FBS for 24 hours and harvested by trypsinization. Lysis and kinase assay were performed according to manufacturer’s instructions (Cell Signaling Technology (CST)). Briefly, cells were lysed using 1x Cell Lysis buffer (CST #9803) supplemented with 1mM PMSF and 1mM Na_3_VO_4_ and ALK-or ROS1-fusion were pulled down described for affinity purification. Washed beads were resuspended in kinase buffer (CST #9802) supplemented with 200µM ATP (CST #9804) and either 1µg recombinant SHP2 (R&D #1894-SH-100) or recombinant STAT3 (Active Motif #81095), respectively, and incubated for 30min at 30°C on a shaker. Reaction was terminated by adding 20µl 4x SDS sample buffer (BioRad 1610747). Analysis of assay was performed by loading equal volumes for Western blotting.

### Western blot

Whole cell lysates were prepared using RIPA buffer as described above and boiled containing 1x SDS sample buffer and equal protein concentrations were loaded. Immunoprecipitation was performed as described above and beads were boiled in 2x SDS sample buffer and equal volumes were loaded. Cellular fractionation was performed using a commercial product and following the manufacturers protocol (Thermo Fisher #78840), protein concentrations were established using BCA assay, boiled in 1x SDS sample buffer and equal protein concentrations were loaded. Samples, and protein ladder (Thermo Fisher #26619) were loaded on tris-glycine 4%-20% gradient gels (BioRad # 4561094) and run in tris-glycine buffer. Transfer on 0.2µm nitrocellulose membranes (BioRad # 1704158) using a semi-dry transfer device (BioRad # 1704150) using preset settings. Membranes were washed once in 1xTBS and blocked in 5% milk in 1xTBST (TBS + 0.1% Tween20) for 1hour at room temperature. After 1 wash in 1xTBST membranes were incubated in primary antibody solutions, either in 5%milk in 1xTBST for total proteins or 5%BSA in 1xTBST for phospho-proteins overnight at 4°C. After 3 washes with 1xTBST membranes were incubated in HRP-coupled secondary antibody diluted 1:5000 in 5%milk in 1xTBST at room temperature for 1 hour. After 3 washes in 1xTBST and 1 wash in 1xTBS membranes were developed using an automated imaging system (BioRad ChemiDoc Imaging System) and ECL substrates (abundant proteins: BioRad #1705061, scarce proteins: Thermo Fisher #34095) using preset settings for optimal exposure. Secondary antibody stripping was performed by incubating membranes for 15min on shaker in buffer solution (Thermo Fisher #21063) followed by 3 washes in TBST and re-blocking in 5% milk in TBST. Reprobing was only done for antibodies of different host species. Antibodies used in this study: mouse-α-ALK (CST #3791), rabbit-α-p-ALK(Y1507) (CST #14678), mouse-α-ROS1 (CST #3266), rabbit-α-p-ROS1(Y2274) (CST #3078), mouse-α-SHP2 (Abcam #ab76285), rabbit-α-p-SHP2(Y580) (CST #5431), mouse-α-MEK1/2 (CST #4694), rabbit-α-p-MEK1/2 (S217/221) (CST #9154), mouse-α-ERK1/2 (CST #4696), rabbit-α-p-ERK1/2(T202,Y204), rabbit-α-GAB1 (CST #3232), rabbit-α-p-GAB1(Y627) (CST #3233), mouse-α-STAT3 (CST #9139), rabbit-α-p-STAT3(Y705) (CST #9145), rabbit-α-p-STAT3(Y705) (*in vitro* kinase only; Abcam #ab267373), mouse-α-STAT1 (CST #9176), rabbit-α-p-STAT1(Y701) (CST #9167), rabbit-α-SHC1 (CST #2432), rabbit-α-p-SHC1(Y239/240) (CST #2434), rabbit-α-SHC3 (Proteintech #12436-1-AP), mouse-α-GAPDH-HRP (Proteintech #HRP-60004), rabbit-α-β-TUBULIN-HRP (CST #5346), anti-rabbit IgG-HRP (CST #7074), and anti-mouse IgG-HRP (CST #7076).

### Spheroid Invasion Assay

Spheroid invasion assays (SIA) were conducted following the protocol described previously (58). Briefly, iNHAs were seeded into low-attachment, U-bottom 96-well microplates (Corning Costar #7007) at a density of 2,500 cells/100µl per well. Spheroids were allowed to form over 24 hours at 37°C in 5% CO_2_. Subsequently, 70 µl of media was replaced with a collagen matrix consisting of 5% sodium bicarbonate [stock 7.5%] (Thermo Fisher # 25080094), 10% 10×DMEM low glucose (Sigma Aldrich #D2429), and 85% PureCol® Type I Collagen Solution (Bovine) [stock 3mg/ml] (Advanced BioMatrix #5005). After 2 hours of polymerization, it was overlaid with 100 µl iNHA growth medium. Spheroids were allowed to invade the collagen matrix over 24 hours and were afterwards stained with Hoechst 33342 (Thermo Fisher #62249) 1:500 in PBS, 3 to 4 hours prior microscopy quantification. Imaging was performed using the Harmony software on a PerkinElmer Operetta microscope.

### Immunofluorescence

5000 iNHA cells were seeded in regular medium in treated (ibiTreat) 8-well chamber microscopy slides (ibidi #80862) and grown for 24 hours. Medium was aspirated and cells were washed once in PBS followed by 15min fixation in 4% formaldehyde in PBS (Thermo Fisher #28906) at room temperature. Following 3 washes with PBS, cells were permeabilized using PBS + 0.5% Tritox-X100 for 15min. After 3 washes in PBS + 0.1% Tween-20 (PBST) cells were blocked for 45min at room temperature using 5% normal donkey serum (Jackson Immuno Research # AB_2337258) in PBST. Incubation with following primary antibodies in blocking solution: rabbit-α-DYKDDDDK-tag (binds same epitope as Sigma-Aldrich α-FLAG M2; 1:500; CST #14793), rabbit-α-ROS1 (1:50; CST # 3287), and mouse-α-β-TUBULIN (1:300; Sigma Aldrich #T5201), overnight at 4°C. After 3 washes with PBST, cells were incubated with secondary antibodies, donkey-α-rabbit-488 (Jackson Immuno Research #711-545-152), and donkey-α-mouse-Cy3 (Jackson Immuno Research # 715-165-150) diluted 1:1000 in blocking solution, for 1 hour at room temperature, protected from light. Cells were washed 3 times with PBST, nuclei were counterstained using 1µg/ml DAPI in PBS (CST #4083) for 10min, washed once more with PBS, and mounted using mounting medium (Vectashield #H-1000). Images were acquired using a Nikon Ti2 inverted microscope, equipped with a 60X/1.2 NA water objective and fluorescence filter sets for GFP and Cy3 and postprocessed using ImageJ.

### Indirect iterative immuno fluorescence – details

Staining and analyses, including UMAP projections, of FFPE embedded brain slices were performed by Apricot Therapeutics according to Kramer *et al*. (40). Primary antibodies used: rabbit-α-ALK (1:60, CST #3633), rabbit-α-ROS1 (1:80, CST #63452), rabbit-α-SHP2 (1:40, CST #3397), rabbit-α-p-SHP2(Y580) (1:60, Thermo Fisher #PA5-114642), rabbit-α-ERK1/2 (1:150, CST #4695), rabbit-α-p-ERK1/2(T202, Y204) (1:80, CST #4370), mouse-α-STAT3 (1:40, CST #9139), rabbit-α-p-STAT3(Y705) (1:60, CST #9145), rabbit-α-ACTIN (1:300, Abcam #ab8227), mouse-α-LaminB1 (1:80, Biolegend #869802), rat-α-KI67 (1:200, Thermo Fisher #17-5698-82), rabbit-α-H3K27me3 (1:40, Abcam #ab192985), rabbit-α-H3K27ac (1:40, Abcam #ab4729), rabbit-α-p-AKT(S437) (1:80, CST #4060), rat-α-SOX2 (1:80, Thermo Fisher # 14-9811-82), mouse-α-NESTIN (1:20, BD-Biosciences #556309), goat-α-SOX10 (1:80, R&D #AF2864), chicken-α-GFAP (1:1500, NovusBio #NBP1-05198), goat-α-OLIG2 (1:20, R&D #AF2418), goat-α-PDGFRα (1:200, R&D #AF1062), rabbit-α-S100β (1:1300, Abcam #ab52642), and rat-α-MBP (1:1500, Abcam #ab7349). Secondary antibodies used,: donkey-α-rabbit IgG (H+L) highly cross adsorbed Alexa Fluor 647 (1:500, Invitrogen #A31573), donkey-α-rabbit IgG (H+L) highly cross adsorbed Alexa Fluor 568 (1:500, Invitrogen #A10042), donkey-α-mouse IgG (H+L) highly cross adsorbed Alexa Fluor 488 (1:500, Invitrogen #A21202), donkey-α-rat IgG (H+L) highly cross adsorbed Alexa Fluor Plus 647 (1:500, Invitrogen #A48272), donkey-α-goat IgG (H+L) highly cross adsorbed Alexa Fluor 647 (1:500, Invitrogen #A21447), and donkey-α-chicken IgG (H+L) highly cross adsorbed Alexa Fluor 568 (1:500, Invitrogen #A78950). Blocking for mouse antibodies was performed using Fab Fragment donkey-α-mouse IgG (H+L) (1:43.3, Jackson Immuno Research #715-007-003).

### Data visualization

ggplot2 (59), ggpubr (60), ComplexHeatmap (46) and EnhancedVolcano (61) R packages were used for the visualization of the boxplots (including statistics), heatmaps and Volcano plots.

### Statistics

Statistical analyses were performed using GraphPad Prism (v10.0.2). Two group comparisons were done using two tailed unpaired Student’s t-test. Multi group comparisons were tested for normal distribution, one-way ANOVAs, parametric for normal distributed data and non-parametric (Kruskal-Wallis) for not-normal distributed data, followed by post-hoc pairwise comparison using Dunn’s test, were performed to establish significance. Correlations were determined using Spearman’s correlation as data are not normally distributed. Survival analysis was performed using the Kaplan-Meier method, with differences in survival being assessed using Mantel-Cox log-rank test.

---

**Figure S1:**
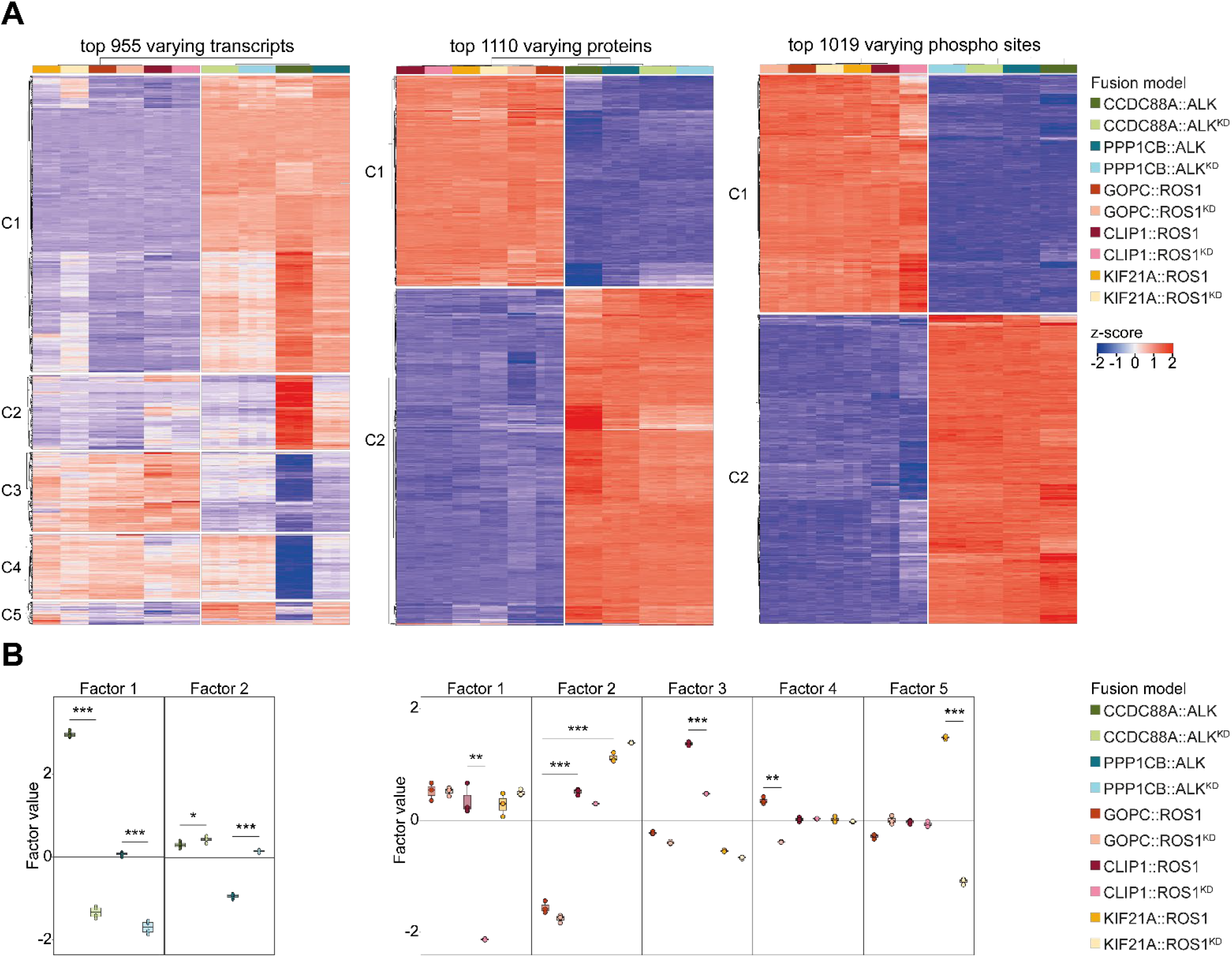
Omic approaches reveal distinct alterations between ALK- and ROS1-fusions as well as between investigated omic layers. **(A)** Heatmaps illustrating top varying molecules detected, color gradient: z-score high (red) to low (blue), samples indicated at the top, clustering based on ALK vs ROS1, (left) top 955 varying transcripts, (middle) top 1110 varying proteins, (right) top 1019 varying phosphosites. **(B)** Factor box plot illustrating the distribution of MOFA factors scores across samples; Factors 1 and 3 observed variance for CLIP1::ROS1, factor 1: transcriptome, factor 3:transcriptome and phospho-proteome, factors 4 and 5 observed variances between GOPC::ROS1 or KIF21A::ROS1, respectively, GOPC1::ROS1: proteome, KIF21A::ROS1:transcriptome; y-axis: factor score, arbitrary units; Wilcoxon rank-sum test, **: p-value <0.01, ***p-value <0.001.

**Figure S2:**
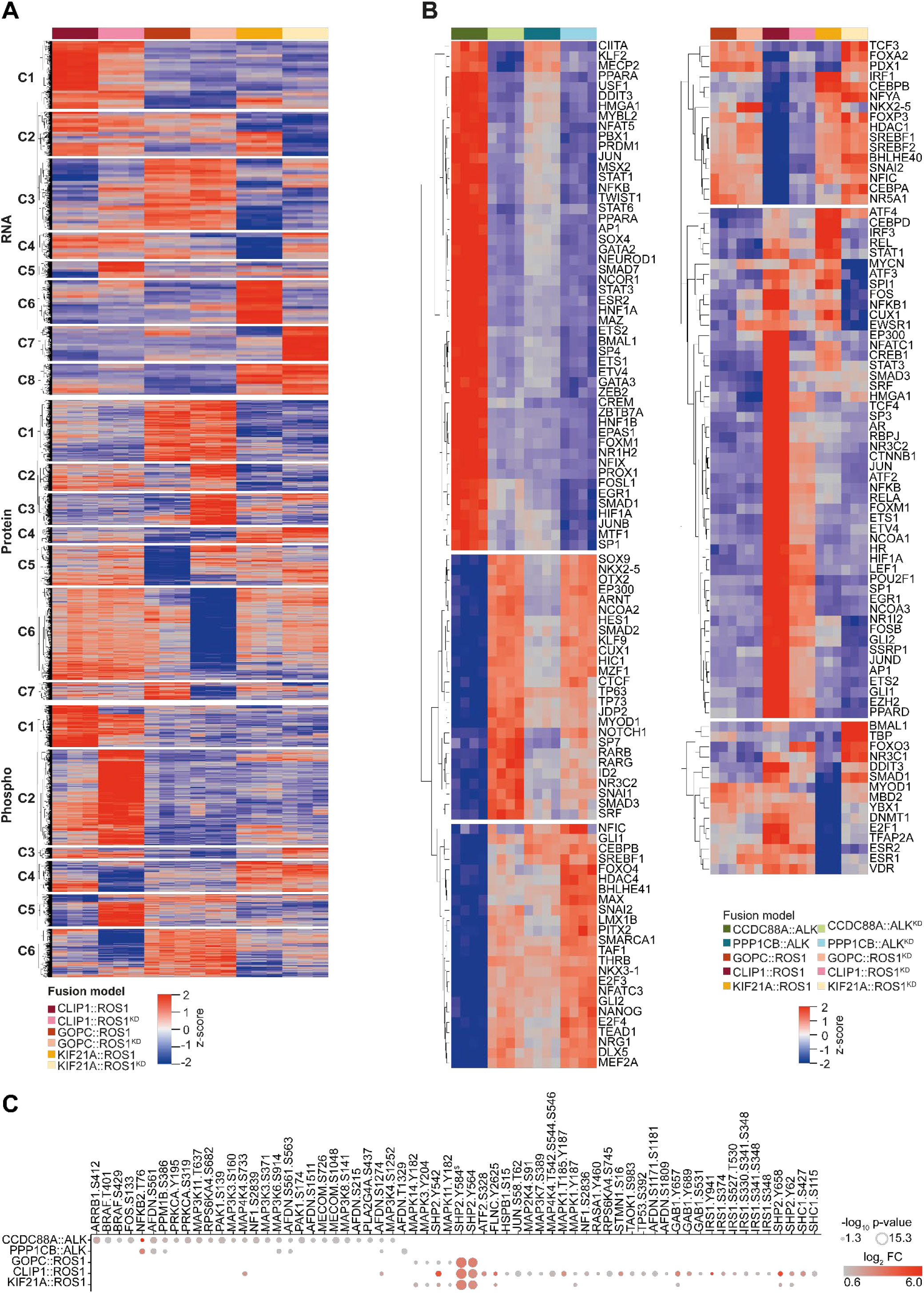
Integrative phosphoproteogenomic analyzes – complete visualizations. **(A)** complete ROS1-fusion MOFA heatmap, color gradient: z-score high (red) to low (blue); related to Fig3E. **(B)** Complete transcription factor activity heatmap for (left) ALK- and (right) ROS1-fusions, color gradient: z-score high (red) to low (blue); related to Fig3F,G. **(C)** Differentially phosphorylated MAPK pathway proteins compared to corresponding KD controls, size: −log_10_ p-value, color gradient: log_2_ FC high (red) to low (grey), ^$^: mass spectrometry identified phosphorylation at Y584, whose site-localization probability was low. Follow-up validation confirmed Y580 as the true phosphorylation site.

**Figure S3:**
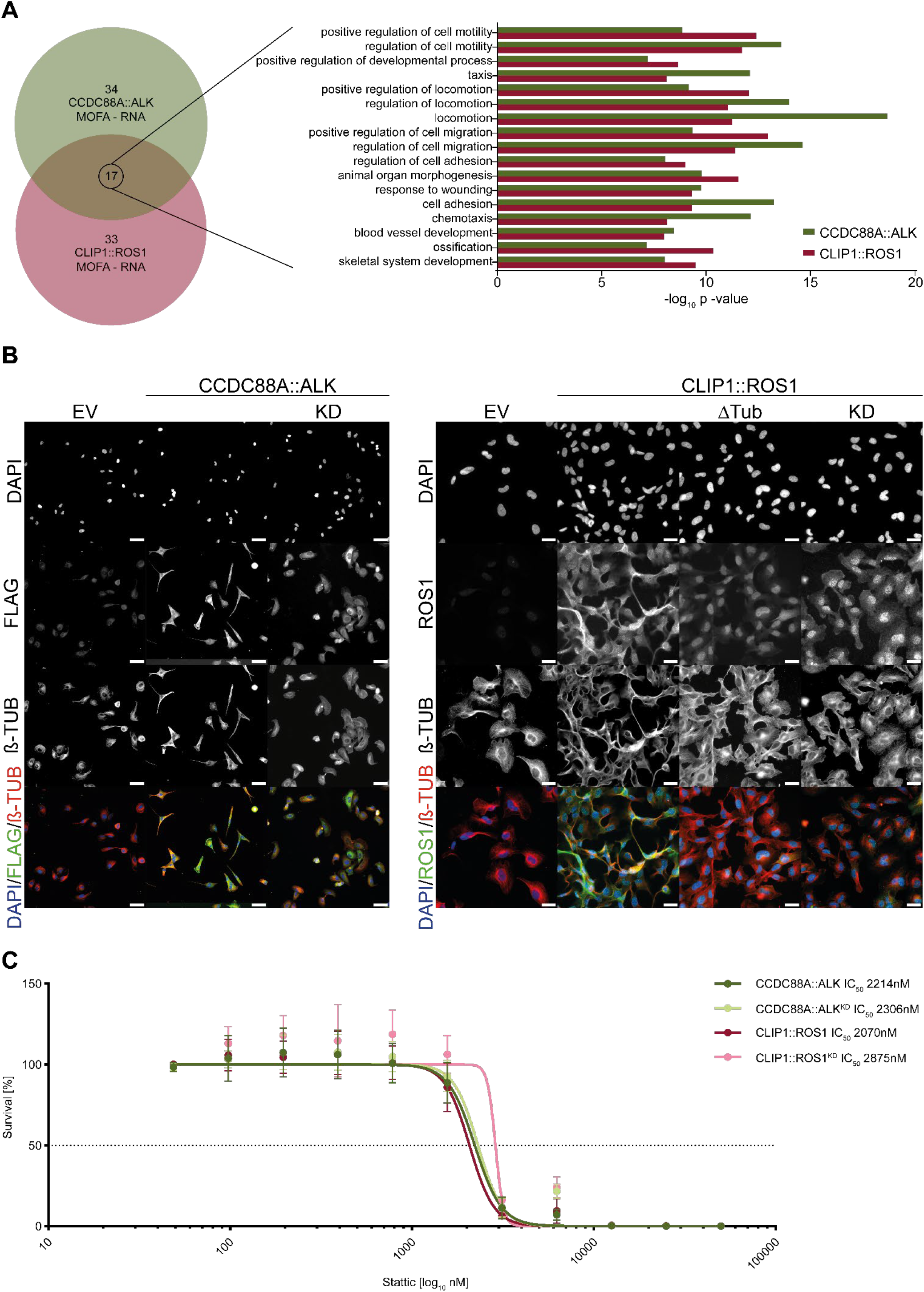
CCDC88A::ALK and CLIP::ROS1 expression result in hyper motile phenotypes presumably facilitated via STAT3 signaling. **(A)** Venn diagram highlighting overlapping biological process GO: terms derived from MOFA weighted transcripts for CCDC88A::ALK (green) and CLIP1::ROS1 (berry), respectively; right: bar plot view on overlapping GO:terms from (A), x-axis: −log10 p-value based on toppgene analysis; y-axis: identified GO:terms, bold: pathways involved in motility; green: CCDC88A::ALK, berry: CLIP1::ROS1. **(B)** Widefield immunofluorescence showcasing overlapping staining of CCDC::ALK or CLIP1::ROS1, respectively, with microtubules; EV: empty vector, KD: kinase dead control; ΔTUB: abrogated microtubule interaction domain; β-TUB: marker for microtubules; scalebar: 10µm. **(C)** Drug titration curves highlighting dose dependent effects of 72h treatment with STAT3i (Stattic) on iNHA models, dark green: CCDC88A::ALK, light green: CCDC88A::ALK^KD^, berry: CLIP1::ROS1; light berry: CLIP1::ROS1^KD^, y-axis: log_10_ drug concentrations, y-axis: survival normalized to DMSO control; dashed line: 50% survival; error bars: SD of 3 biological replicates.

**Figure S4:**
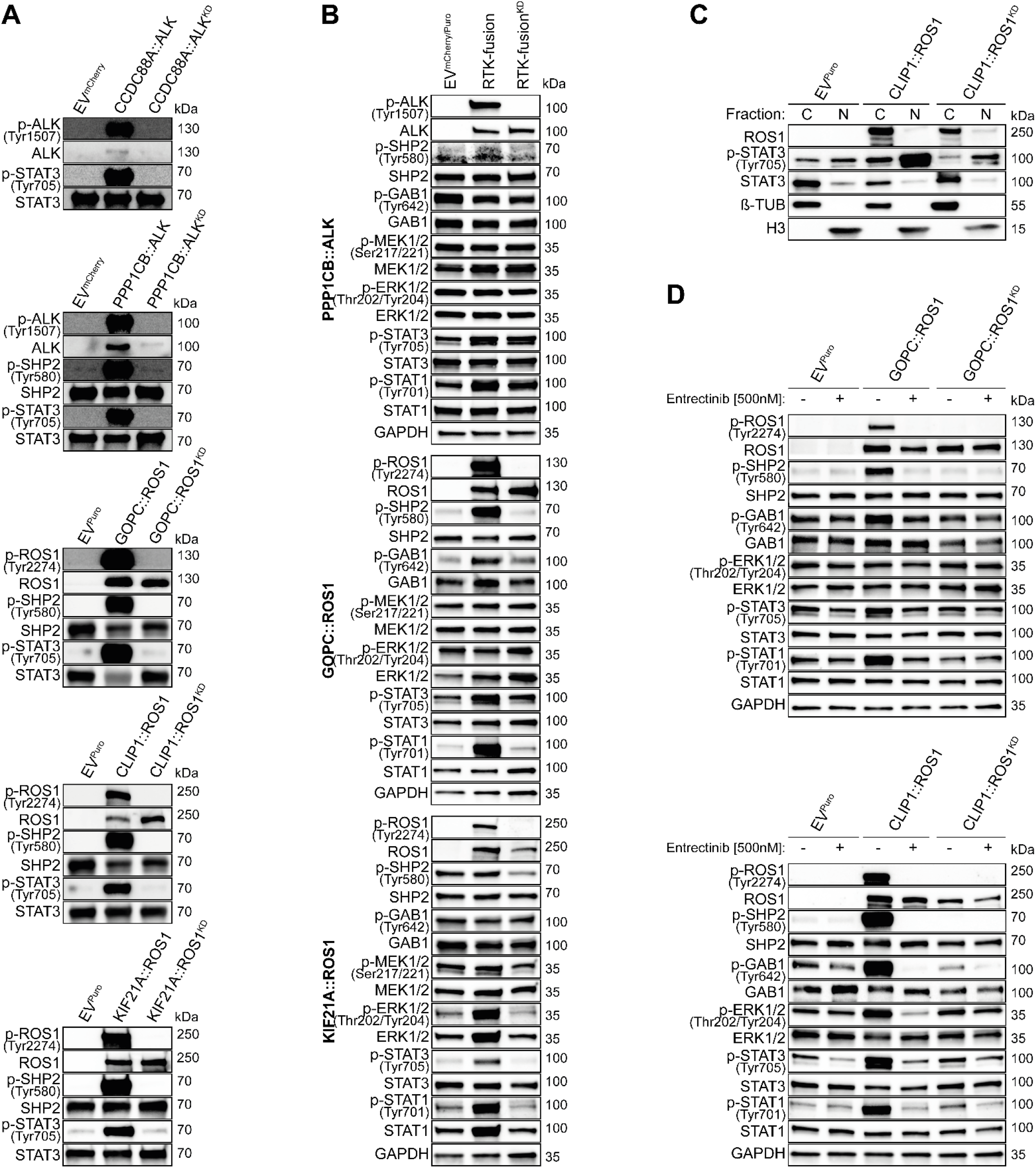
ALK- and ROS1-fusions activate STAT3 and SHP2 through phosphorylation. **(A)** *In vitro* kinase assay validating SHP2 and STAT3 as substrates of ALK- and ROS1-fusions; GAPDH: loading control, phospho-ALK,-ROS1 antibody used to validate KD mutants, phospho-SHP2 (Tyr580) or phospho-STAT3 (Tyr705) validate ALK- and ROS1-fusion kinase specificity towards SHP2 or STAT3, respectively. **(B)** Western blot analysis of MAPK signaling in ALK- and ROS1-fusion models. GAPDH: loading control, phospho-ALK (Tyr1507), -ROS1 (Tyr2274) antibody used to validate KD mutants, phospho-SHP2 (Tyr580), phospho-GAB1 (Tyr642), phospho-MEK1/2 (Ser217/221), and phospho-ERK1/2 (Thr202/Tyr204) used to validate MAPK pathway activity, phospho-STAT3 (Tyr705), and phospho-STAT1 (Tyr701) used to validate STAT activation. **(C)** Subcellular fractionation of CLIP1::ROS1 samples validating increased STAT3 activity; phospho-ROS1 (Tyr2274) antibody used to validate KD mutant and phospho-STAT3 (Tyr705) used to validate pathway activity, β-TUB: cytoplasmic marker, H3: nuclear marker. **(D)** Western blots analyzing the effect of RTK inhibition (Entrectinib 500nM, 4hours) on MAPK and STAT signaling in ALK- and ROS1-fusion models. GAPDH: loading control, phospho-ALK (Tyr1507), -ROS1 (Tyr2274) antibody used to validate inhibition, phospho-SHP2 (Tyr580), phospho-GAB1 (Tyr642), phospho-MEK1/2 (Ser217/221), and phospho-ERK1/2 (Thr202/Tyr204) used to validate MAPK pathway inhibition and phospho-STAT3 (Tyr705), and phospho-STAT1 (Tyr701) for STAT inhibition.

**Figure S5:**
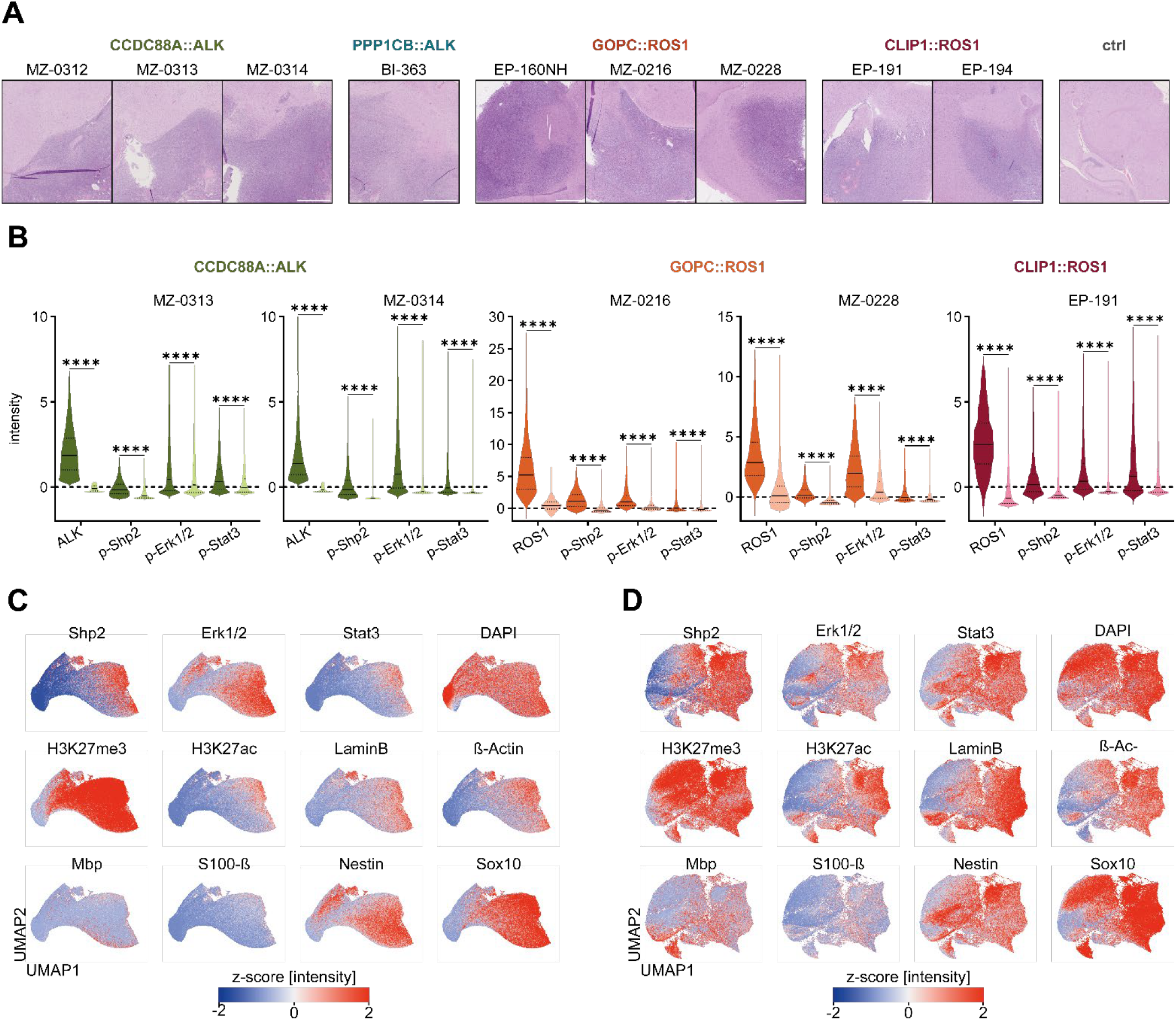
Spatial profiling reveals upregulated Stat3 and Shp2/Mapk signaling in primary ALK- and ROS1 murine tumors. **(A)** H&E stains of brain tumor areas used for multiplex IF, scalebar: 500µm. **(B)** Single cell intensity measurements for indicated antibodies detected by multiplex IF; x-axis: probed proteins, y-axis: z-score normalized intensities; MZ-0313, MZ-0314: CCDC88A::ALK tumors; MZ-0216, MZ-0228: GOPC::ROS1 tumors, EP-194: CLIP1::ROS1 tumor; x-axis: probed proteins, y-axis: z-score normalized intensities; visualization: violin plot, indicating interquartile range and median, Kruskal-Wallis-test, post-hoc Dunn-Bonferroni test: ****:p-value<0.0001; related to Fig5B. **(C-D)** UMAP representation of marker intensities at single cell resolution, normalized to healthy control; color gradient: normalized z-score high (red) to low (blue); **(C)** ALK-samples, **(D)** ROS1-samples.

**Figure S6:**
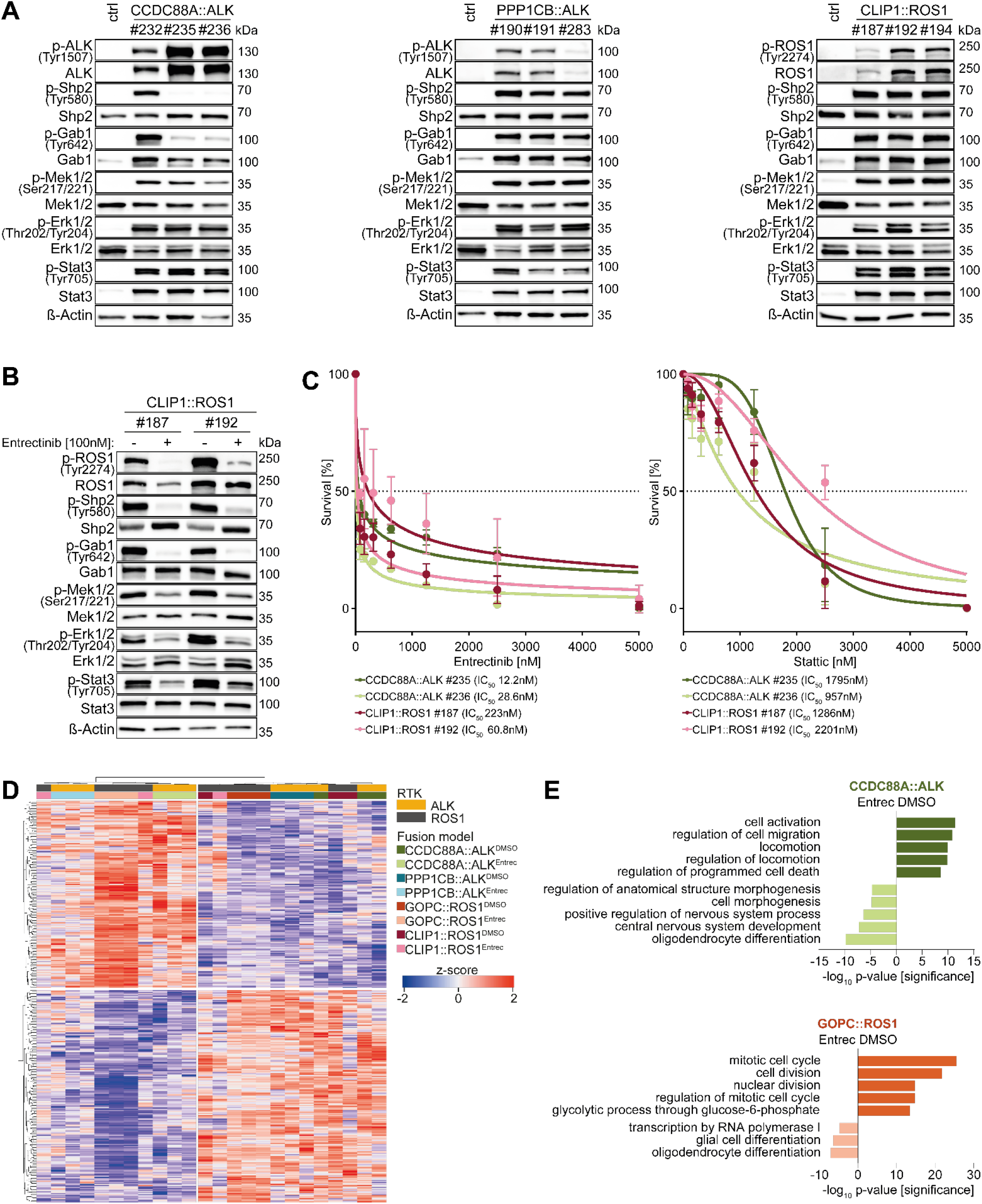
Murine brain tumor derived cells recapitulate upregulated Stat3 and Shp2/Mapk signaling. **(A)** Western blot analysis of ALK- and ROS1-fusion expression and Shp2/Mapk and Stat3 signaling in ALK- and ROS1-fusion IUE models. β-Actin: loading control, phospho-ALK (Tyr1507), -ROS1 (Tyr2274) antibody used to validate fusion transgene activity, phospho-Shp2 (Tyr580), phospho-Gab1 (Tyr642), phospho-Mek1/2 (Ser217/221), and phospho-Erk1/2 (Thr202/Tyr204) used to validate Mapk pathway activity and phospho-Stat3 (Tyr705) for Stat3 activation. **(B)** Western blots analyzing the effect of RTK inhibition (Entrectinib 100nM, 24hours) on Mapk and Stat3 signaling CLIP1::ROS1-fusion IUE models. β-Tub: loading control, phospho-ROS1 (Tyr2274) antibody used to validate inhibition, phospho-Shp2 (Tyr580), phospho-Gab1 (Tyr642), phospho-Mek1/2 (Ser217/221), and phospho-Erk1/2 (Thr202/Tyr204) used to validate Mapk pathway inhibition and phospho-Stat3 (Tyr705) for Stat3 inhibition. **(C)** Drug titration curves highlighting dose dependent effects of 72h treatment with Entrectinib (left), or Stattic (right) on IUE models, dark green: CCDC88A::ALK #235, light green: CCDC88A::ALK #236, berry: CLIP1::ROS1 #187; light berry: CLIP1::ROS1 #192, y-axis: linear drug concentrations, y-axis: survival normalized to DMSO control; dashed line: 50% survival; error bars: SD of 3 biological replicates. (**D)** Unsupervised clustering, Euclidian distance with complete linkage heatmap of most variable transcripts, color gradient: z-score high (red) to low (blue), samples indicated at the top. **(E)** Enriched GO:terms for DEG in CCDC88A::ALK (upper) or GOPC::ROS1 (lower) samples. x-axis: −log_10_ p-value significance established by ReViGo; left: GO:terms enriched in Entrectinib treated samples, right: GO:terms enriched in DMSO control samples

